# Trait stability of diverse kabuli chickpea germplasm from delayed sowing in a rainfed environment

**DOI:** 10.64898/2026.05.29.728723

**Authors:** Callum B. Jamie, Shanice Van Haeften, Victor Papin, Alison Kelly, Karine Chenu, Jingyang Tong, Cara Jeffrey, Laura Ziems, Lee T. Hickey, Richard Trethowan, Millicent R. Smith

**Affiliations:** Centre for Crop Science, Queensland Alliance for Agriculture and Food Innovation, The University of Queensland, Brisbane, Queensland, Australia; ARC Training Centre in Predictive Breeding, The University of Queensland, St Lucia, QLD 4072, Australia; ARC Hub for Engineering Plants to Replace Fossil Carbon, The University of Queensland, St Lucia, QLD 4072, Australia; The Plant Breeding Institute, The University of Sydney, Sydney, NSW, Australia; School of Agriculture and Food Sustainability, Faculty of Science, The University of Queensland, Queensland, Australia

**Author notes:** **Correspondence:** Richard Trethowan and Millicent R. Smith are co-corresponding authors.; Richard Trethowan,; Millicent Smith.

**Keywords:** *Cicer arietinum*, critical period, delayed sowing, haplotype, heat stress, high temperature

## Abstract

**Context and Objective:** Delayed sowing can expose chickpea crops to stress during the critical period for yield determination, but the effect of yield components and phenology to grain yield variation is not well characterised in diverse germplasm under rainfed conditions. Identifying genetic resources for grain yield improvement requires integration of multi-environment trial and genomic analyses to disentangle direct yield effects from indirect effects of phenology. This study aimed to (1) characterise genotype by environment interaction patterns for grain yield, yield components and phenology across times of sowing and seasons, and (2) identify genomic regions associated with improved grain yield that are present in genebank accessions but absent from current Australian commercial cultivars.

**Methods:** A diversity panel of 141 kabuli chickpea genotypes, including six commercial Australian cultivars and 135 genebank accessions, was evaluated across six rainfed trials at Narrabri, New South Wales over three seasons (2018 to 2020) under typical (MAIN) and delayed (LATE) sowing. Multi-environment trial analyses with factor analytic models partitioned genotype by environment interactions for grain yield, 100-seed weight, seed number, and thermal time to flowering, podding, and maturity. Haplotype block analysis identified high variance blocks associated with seed number, classified by their overlap with high variance thermal time to flowering blocks, to distinguish from effects mediated by phenology.

**Results and Conclusions:** Delayed sowing reduced grain yield by up to 1.04 t ha^-^¹, driven primarily by reductions in seed number rather than 100-seed weight. Accelerated phenology was a key component of adaptation among commercial cultivars. Four haploblocks with high block variance for seed number were identified across all six trials.

**Significance:** Seed number was the dominant driver of grain yield variation in this diverse kabuli chickpea panel. Targeted introgression of rare superior haplotypes from genebank accessions provides an opportunity to broaden the genetic base of Australian kabuli chickpea and improve yield through higher seed number, with relevance to chickpea production systems facing similar climate variability.

**Highlights:** Delayed sowing reduced grain yield in diverse kabuli chickpea germplasm by up to 1.04 t ha^-1^ across three years and six trials in northern New South Wales.

Seed number, not seed weight, was the dominant driver of grain yield variation, and a shorter phenological duration was associated with higher seed number.

Across all six trials, haplotype block analysis identified four genomic regions in high linkage disequilibrium with high variance for seed number and low variance for flowering time.

The accession FLIP 94 62C uniquely carried rare superior haplotypes at two chromosome 4 blocks, the haplotype at the 17.0 Mb block was the most superior haplotype in all trials while the haplotype at the 8.5Mb block was most superior only in the most heat stressed environment.

## 1. Introduction

Chickpea (*Cicer arietinum* L.) is an annual grain legume valued as an affordable source of protein, carbohydrates, vitamins and micronutrients. Given malnutrition and micronutrient deficiencies affect over 2 billion people worldwide, chickpea production offers a means to address nutritional insecurity in subsistence systems that staple cereals alone cannot satisfy (Jha et al., 2024; Weffort & Lamounier, 2024). In Australia, interest in chickpea production has risen through demand for plant based protein and its suitability in cereal based crop rotations as it improves soil fertility through nitrogen fixation and disrupts pest and disease cycles (Berger et al., 2004; Merga & Haji, 2019). Despite growing importance, productivity remains low and unstable (Anwar et al., 2022) with an average grain yield of 1.17 tonnes per hectare over 14.0 million hectares worldwide (FAO, 2023). Yield limitation is largely driven by abiotic stress, which can cause more than 70% yield loss (Varshney et al., 2019), and narrow genetic diversity in the elite gene pool has impeded success of conventional breeding approaches to develop superior genotypes with stress adaptation (Varshney et al., 2009).

Growing degree days (°Cd) are used to quantify accumulated heat units and describe the timing of phenological development (McMaster & Wilhelm, 1997). Extensions to this framework have been used in chickpea to quantify cumulative exposure to temperatures above the optimum, and revealed negative associations of heat with both grain yield and seed size (Jeffrey et al., 2025). This prior characterisation of the germplasm panel used in the present study, evaluated at Narrabri, NSW and Kununurra, WA, demonstrated that growing degree day models can effectively quantify temperature stress responses across field environments. Chickpea is most sensitive to stress during the ‘critical period’ of yield development, which spans 800°Cd, centred 100°Cd after flowering. During this period, growth rate is correlated linearly to yield (Lake & Sadras, 2014, 2016). Heat stress has emerged as a major yield limitation because chickpea is widely cultivated in systems that expose it to rising temperatures during this period, such as in post-rainy season subtropical systems or winter-sown Mediterranean environments (Davies et al., 1999; Lake & Sadras, 2014). In Australia’s major growing regions of northern New South Wales and southern Queensland, chickpea production is particularly exposed as sowing can be delayed by a lack of early rainfall, wet weather, or through deliberate management to reduce incidence of the devastating fungal disease, Ascochyta blight (Devasirvatham et al., 2012). High temperatures during reproduction can cause flower and pod abortion, reduced pollen viability, and impaired grain filling which leads to yield loss (Devasirvatham et al., 2015b; Kaushal et al., 2013). Climate change is predicted to increase both the mean temperature and the frequency of acute high temperature events which will exacerbate heat stress and yield instability (Lake et al., 2025; Masson-Delmotte et al., 2021), highlighting the need for superior germplasm adapted to these environments.

Developing heat adapted cultivars remains a challenge as heat tolerance is expressed through complex responses that vary dynamically with the timing, intensity, and duration of stress exposure during developmental stages (Jha et al., 2025; Rani et al., 2020). Yield and adaptive traits that confer tolerance are typically polygenic and characterised by low heritability and large genotype by environment interactions (GEI) (Arriagada et al., 2022; Danakumara et al., 2024). This GEI complicates identification of superior germplasm for progression in breeding programs, particularly when they exhibit crossover patterns and genotype rankings change across environments (Baker, 1988; Stella et al., 2025).

Delayed sowing trials are commonly used as a practical and inexpensive method to screen large numbers of genotypes to elevated temperatures in field conditions (Devasirvatham et al., 2015a; Hiremath et al., 2011), however comparisons are confounded by changes in environment such as photoperiod, radiation, vapor pressure deficit and rainfall (Sadras et al., 2015). Recent advances in multi-environment trial (MET) analysis and environmental characterisation have improved the capacity to disentangle direct temperature effects from the indirect effects, including phenological shifts caused by temperature dependent development rates (Bonhomme, 2000; Sadras et al., 2015). To identify effective breeding targets under heat stress, it is critical to determine which yield components drive GEI and separate direct component effects from the indirect effects of phenology.

Improved methods to integrating genomic and phenotypic data enables breeders to quantify GEI, understand the genetic architecture of adaptation, and identify germplasm with specific or broad adaptation across the TPE (Piepho & Blancon, 2023; Shaffer et al., 2025; Smith & Cullis, 2018; Voss-Fels et al., 2019). Delayed sowing studies in chickpea have predominantly evaluated phenotypic responses without genomic characterisation (Devasirvatham et al., 2015a; Krishnamurthy et al., 2011), and while genomic approaches have identified quantitative trait loci associated with heat and drought adaptation (Barmukh et al., 2022; Garg et al., 2025; Jeffrey et al., 2024; Varshney et al., 2014), including heat tolerance QTL identified in this germplasm panel studied at Narrabri, NSW and Kununurra, WA (Jeffrey et al., 2024), the integration of MET frameworks that explicitly model GEI in rainfed production environments is limited.

Building on the prior characterisation of this germplasm panel (Jeffrey et al., 2024, 2025), we apply a MET framework with factor analytic models to formally partition GEI for yield and its component traits across six rainfed trials at Narrabri, NSW over three seasons from 2018 to 2020 under typical (MAIN) and delayed (LATE) sowing. We focus on kabuli germplasm evaluated in Narrabri only to avoid confounded effects of seed type on seed weight and photoperiod differences across locations. Specifically, this study aims to (1) characterise the relationships between grain yield, yield components (seed number and seed weight), and phenology across environments generated by differences in time of sowing and season, (2) identify genotypes that exhibit either specific adaptation to high temperature conditions or broad adaptation across environments, and (3) identify genomic regions and haplotype blocks associated with seed number that are independent of phenology effects, to locate superior haplotypes absent from elite Australian cultivars.

## 2. Materials and Methods

### 2.1. Plant materials

A diversity panel of 157 genotypes was evaluated in this study, which included 143 kabuli- and 14 desi-type chickpea sourced from the Australian Grains Genebank (AGG), the International Center for Agricultural Research in the Dry Areas (ICARDA) and the International Crops Research Institute for the Semi-Arid Tropics (ICRISAT) (Supplementary Table 1). The panel was comprised of germplasm accessions predominantly originating from Syria but also included accessions from India and released cultivars from Australia. This panel was assembled based on prior testing by ICARDA and ICRISAT (determined to be high yielding in response to heat stress) and to capture both landrace diversity and improved germplasm (Figure 1). A minimum of 145 genotypes were concurrent across all trials.

**Figure 1.**
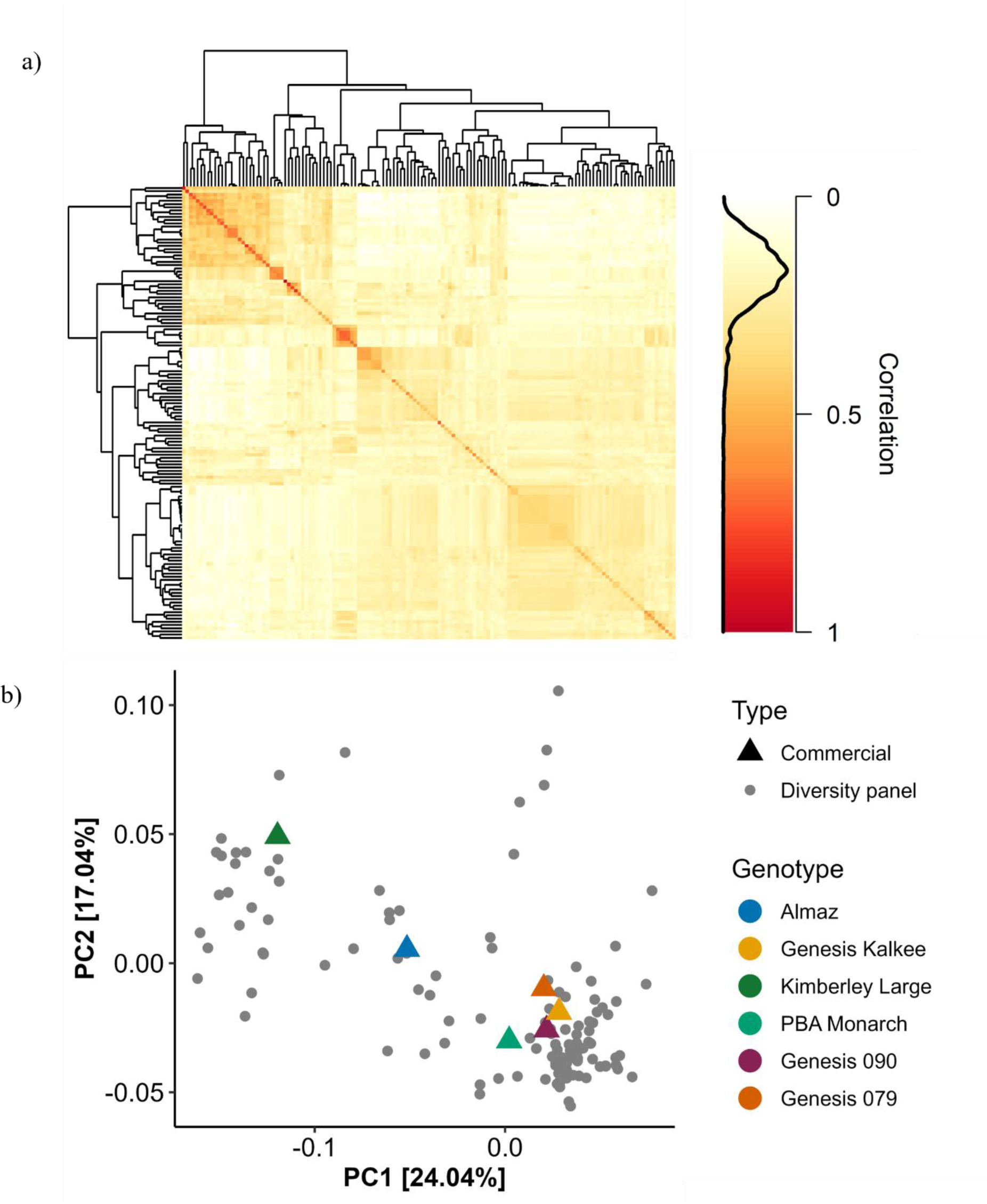
Genomic relationships and population structure of the chickpea diversity panel. a) Heatmap of the genomic relationship matrix (GRM) showing genetic similarity among genotypes (white = 0, red = 1), with hierarchical clustering displayed in the dendrogram. b) Principal coordinate analysis (PCoA) based on 8,093 SNP markers showing population structure, with commercial (triangles, coloured) and diversity panel (circles, grey) genotypes distinguished by shape and colour.

Out of the genotypes evaluated in the field trials, 155 were genotyped with an Infinium™ Pulses 30K v1.0 bead chip array (Illumina Inc., San Diego, USA) against the CDC Frontier reference genome v1.0 (Varshney et al., 2013) and imputed to high density using MiniMac3 (Das et al., 2016) to a set of 765,703 SNPs. Marker curation was conducted in PLINK v1.9 (Chang et al., 2015; Purcell & Chang, 2025). Markers with a minor allele frequency less than 1% were removed. To account for highly correlated markers, linkage disequilibrium (LD) pruning applied a sliding window of 1,000 SNPs, a step size of 10 SNPs, and an R^2^ threshold of 0.99. This procedure retained one marker from each set of SNPs in high LD and resulted in 8,093 SNP markers retained for downstream analyses. Population structure analysis was performed on kabuli genotypes with the *SelectionTools* package. Genetic distance between genotypes was calculated using Rogers’ distance metric (Rogers, 1972). The first 10 principal components were extracted, and their variance contributions were determined from eigenvalues (Figure 1b).

### 2.2. Field trials

A subset of datasets that were analysed by Jeffrey et al. (2024; 2025) were used in this study. This includes a total of six field trials which were conducted across three years (2018 to 2020) and two times of sowing (TOS). Trials were established at The University of Sydney, Narrabri, New South Wales (30.2737° S, 149.7350° E) which represented a typical Australian production environment in a Mediterranean climate grown under rainfed conditions. Due to exceptionally low seasonal rainfall, both trials in 2019 received supplementary irrigation, including two applications of 40 mm on the 26^th^ and 29^th^ of April before sowing.

The timing of the first TOS (MAIN) followed regional commercial practice (GRDC, 2016), and the delayed TOS (LATE) was designed to expose plants to higher temperatures during reproductive development. The LATE sowing was delayed relative to the MAIN sowing by 41 days in 2018, 57 days in 2019 and 59 days in 2020 (Table 1).

**Table 1.** Summary of sowing dates (DD/MM), time of sowing (TOS) classification, and genotype replication for six trials at Narrabri from 2018 to 2020

| Year | Sowing date | TOS | Number of genotypes | Replicates |  |  |
| --- | --- | --- | --- | --- | --- | --- |
|  |  |  |  | 1 | 2 | >2 |
| 2018 | 21/06 | MAIN | 145 | 67 | 72 | 6 |
|  | 01/08 | LATE | 154 | 27 | 119 | 7 |
| 2019 | 05/06 | MAIN | 151 | 5 | 143 | 3 |
|  | 01/08 | LATE | 151 | 4 | 147 | 0 |
| 2020 | 03/06 | MAIN | 150 | 4 | 147 | 0 |
|  | 01/08 | LATE | 150 | 4 | 147 | 0 |

In 2019 and 2020, genotypes were evaluated in a randomised complete block design while 2018 was an unbalanced design where most genotypes were represented by two or more replications, while some only had one. Plants in all trials were sown in 8 m^2^ plots. Sowing rates followed local commercial standards of approximately 169 seeds per plot (21 plants per m^2^). Trial management followed regional agronomic guidelines (GRDC, 2016). Seed was treated with P-Pickel T (Nufarm) at an application of 2 ml per kg of seed to minimise risk of seed borne disease and were sown at 9 cm depth. Seed was inoculated with TagTeam^®^ peat (Novozymes) at sowing, and a starter fertiliser of Granulock Z Extra (Incitec Pivot Fertiliers) at 80 kg ha^-1^ was applied pre-emergence. Weeds, pests, and diseases were managed as required based on commercial recommendations (GRDC, 2016).

### 2.3. Data collection

#### 2.3.1. Phenology and yield related traits

Days to flowering (DTF), podding (DTP), and maturity (DTM) were recorded as the number of days from sowing until at least half the plants within a plot reached each developmental stage. Flowering was defined as the date at which 50% or more of plants had one or more open flowers, podding was the date at which 50% or more of plants had set one or more pods, and maturity was the date at which 50% or more of plants had brown pods. Plots were harvested with a plot header when the entire experiment achieved harvest maturity. After harvest, grain yield (GY; t ha⁻¹) was recorded by weighing the harvest of each plot, and 100-seed weight (HSW; g) was recorded as a random sample of 100 seeds from each harvest bag. Seed number (SN; seeds m⁻²) was derived for each plot from GY and HSW, where GY was converted to g ha⁻¹, divided by the weight of one seed (HSW / 100), and divided by 10⁴ to give seeds _m_-2.

Chickpea development was quantified with thermal time (Tt) expressed as growing degree days (GDD; °Cd). Daily GDD was calculated using a segmented function to account for reduced development below and above the optimum temperature (Equation 1).

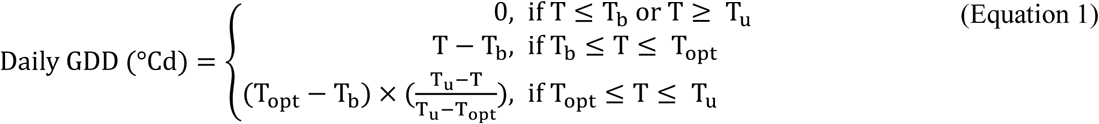

where T is the daily mean temperature, T_b_ is the base temperature (0°C), T_opt_ is the optimum temperature (30°C), and T_u_ is the upper threshold temperature (40°C) (Robertson et al., 2002). Daily GDD values were accumulated throughout the growing season to describe the timing of phenological development (Zhou & Wang, 2018), including DTF, DTP and DTM. Further adjustments for photoperiod and soil moisture (Chauhan et al., 2019; Soltani et al., 2006; Soltani & Sinclair, 2011) were not applied due to germplasm diversity and lack of corresponding data.

#### 2.3.2. Environmental covariates

Weather data was collected from an on-site weather station which sampled every 15 minutes. Recorded variables included daily average (T_avg_; °C), maximum (T_max_; °C), and minimum (T_min_; °C) temperature, rainfall (mm), relative humidity (RH; %), vapour pressure deficit (VPD; kPa), and solar radiation (MJ m^−2^ day^−1^). Daily photoperiod (Pp; hours of daylight) was retrieved from the R package *geosphere* (Hijmans, 2024).

These raw weather variables were transformed to derive biologically relevant covariates. A day over 32°C (DO32) was defined as any day where T_max_ exceeded 32°C. Photosynthetically active radiation (PAR; mol m^−2^ day^−1^) was estimated as incident solar radiation multiplied by 2.1, combining the photosynthetically active fraction of total radiation (∼0.48) and the conversion from energy to photon flux (Meek et al., 1984). Photothermal quotient (PTQ) was calculated for each growth window as the ratio of total incident PAR to the sum of daily mean temperatures over the window (Fischer, 2016; Nix, 1976). An adjusted PTQ incorporated VPD to account for atmospheric demand, and was calculated as the ratio of PTQ to VPD (PTQ_VPD_) (Rodriguez & Sadras, 2007).

### 2.4. Statistical analysis

#### 2.4.1. Environmental covariates

Environmental covariates were calculated for each plot across two growth windows including sowing to flowering (S-F) and during the critical period (Crit). The critical period was defined as 300°Cd before and 500°Cd after flowering (Lake & Sadras, 2014). The flowering date for each plot was derived from Best linear Unbiased Estimates from single site spatial analyses with genotype as a fixed effect, and the calendar dates corresponding to the critical period GDD boundaries were back- and forward-calculated.

#### 2.4.2. Multi-environment trial analysis

Each trait analysed in this study (GY, HSW, SN, DTF, DTP, DTM) was analysed using a multi-environment trial (MET) framework based on linear mixed models (Equation 2) fitted in ASReml-R Version 4.2 (Butler et al., 2023) within R Version 4.5.1 (R Core Team, 2025). This approach accounted for GEI and allowed appropriate modelling of trial-specific error variance structures.

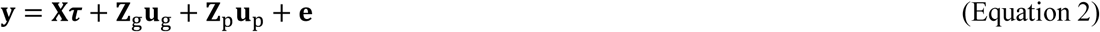

where 𝒚 is the vector of observed responses, 𝐗 is the design matrix for fixed effects and 𝝉 is the vector of fixed effect coefficients, 𝐙_g_ is the design matrix for genetic effects 𝐮_g_, partitioned into additive and non-additive components, 𝐙_p_𝐮_p_ is the design matrix and random effects for trial specific structures, and 𝐞 is the residual error vector. Fixed effects included trial, chickpea seed type (Kabuli, Desi), their interaction, and, where identified in single site spatial analyses, linear row and column trend covariates. Random trial specific effects included blocking and row or range effects where these were significant. Desi genotypes were included as a fixed effect to account for type differences in performance but were excluded from the genomic analysis due to insufficient numbers (n = 14) to reliably estimate GEI.

To account for local spatial trends within each trial, the residual error term (𝐞) was modelled using a separable first order autoregressive process (AR1 × AR1) within each trial (Gilmour et al., 1997). This structure captures correlation between neighbouring plots in both row and column directions, while assuming independence of residuals across trials. Single site spatial analyses were first conducted to identify significant fixed and random spatial effects for each trial. Outliers were detected by inspecting studentised residuals exceeding 4, and removed iteratively where the model was refit after each removal step to check for additional outliers (Piepho et al., 2008). The final model terms retained for each trait and trial are listed in Supplementary Figure 1.

A genomic relationship matrix (GRM) was constructed using the VanRaden (2008) method (1) from 8,093 pruned SNP markers. Genetic effects were partitioned into additive and non-additive components (Equation 3).

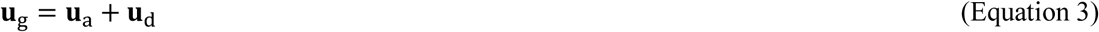

where 𝐮_g_ is a vector of total genetic effects for all genotypes across trials, 𝐮_a_ is the additive genetic effects and 𝐮_d_ is the non-additive genetic effects. Different genetic variance structure models were tested to determine the optimal structure for modelling GEI across trials. Genotypes without marker data were excluded from the genomic model. Broad-sense heritability (H²) was estimated from a diagonal (DIAG) model (Cullis et al., 2006) (Equation 4).

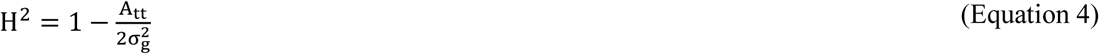

where A_tt_ is the average standard error of the difference between genotype BLUPs and σ^2^ is the genotypic variance. Following, genetic variance-covariance structures of increasing complexity were tested sequentially including a homogeneous correlation model (corv), a heterogeneous correlation model (corh), and factor analytic (FA) models of increasing order. Model selection was based on REML log likelihood and AIC (Smith et al., 2004). After confirmation of heterogeneous variances and genetic correlations across trials, additive genetic effects were modelled using a FA structure combined with the GRM (Equation 5).

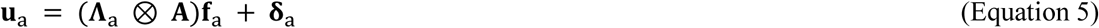

where 𝚲_a_ is an 𝑡 × 𝑘 matrix of the factor loadings across 𝑡 (6) trials, 𝐀 is the additive GRM, 𝐟_a_ is a vector of 𝑘 latent genetic factors, and 𝛅_a_ is a vector of trial specific genetic deviations. The covariance structure of additive genetic effects across trials (Equation 6) was

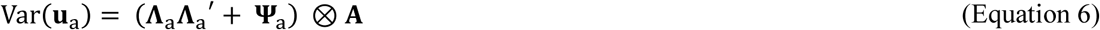

where 𝚲_𝑎_𝚲_𝑎_^′^ captures the genetic covariance structure across trials through shared factor loadings and 𝚿_𝑎_ is a diagonal matrix of trial specific residual additive genetic variances not explained by the common factors (Kirkpatrick & Meyer, 2004). Non-additive genetic effects (𝐮_d_) were modelled using a FA structure with trial specific variances independent of the GRM (Equation 7).

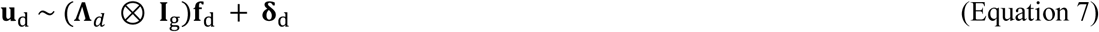

where 𝚲_𝑑_ is an 𝑡 × 𝑘 vector of factor loadings across 𝑡 (6) trials, 𝐈_g_ is an identity matrix of dimensions equal to the number of genotypes (141), 𝐟_d_ is a vector of latent non-additive factors (1 in all traits), and 𝛅_d_ is a vector of trial-specific non-additive deviations. The covariance structure of nonadditive genetic effects (Equation 8) was

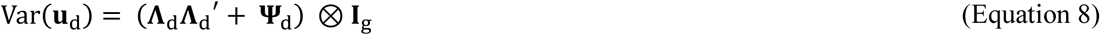

where 𝚿_d_ is a diagonal matrix of trial-specific residual nonadditive variances. This component captured genetic variance not attributable to additive genomic relationships. The number of factors (𝑘) was determined separately for each trait. From the final model, Best Linear Unbiased Predictions (BLUPs) were obtained for each genotype within each trial by predicting over the trial specific factor structure. An additional set of BLUPs marginalised across all six trials were obtained by predicting over the trial main effects, which provided a single across-trials performance estimate per genotype for downstream analyses.

#### 2.4.3. Overall performance and stability

Overall Performance (OP) and Root Mean Square Deviation (RMSD) were derived from the final FA2 model for GY. The unrotated factor loadings were rotated using singular value decomposition (SVD), with genotype factor scores rotated by the same transformation matrix (Smith & Cullis, 2018). OP was calculated for each genotype as the product of its rotated factor 1 score and the sum of rotated factor 1 loadings across trials. RMSD was calculated as the square root of the mean squared trial specific interaction values derived from factor 2, where each genotype’s rotated factor 2 score was multiplied by the corresponding trial’s rotated factor 2 loading. A higher OP indicates better average performance and a lower RMSD indicates greater stability across trials.

#### 2.4.4. Bivariate genetic correlations

Bivariate FA models were used to estimate genetic correlations between trait pairs (GY with SN, HSW, and SN with DTF, DTP, DTM). Each bivariate analysis included two traits across six trials, resulting in 12 trait-trial combinations per analysis. Fixed and random effects for spatial structure were identified in single site spatial analyses. Additive genetic effects were modelled using FA structures with the GRM and non-additive genetic effects were modelled using FA structures with an identity matrix. FA models of increasing order (FA1 to FA3 for additive effects) were tested sequentially and the final model was selected based on likelihood ratio test. Genetic correlations between trait-trial combinations were extracted to quantify trait relationships within each trial.

#### 2.4.5. Haploblock analysis

To identify genomic regions associated with SN independent of phenology, a haplotype-based approach using local genomic estimated breeding values (localGEBVs) was adopted in preference to conventional single marker GWAS. Association signals are often spread across multiple correlated SNPs that are in LD with the same causal variant, reducing power to detect causal genomic regions, particularly in panels of small to moderate size (Shaffer et al., 2025). The localGEBV approach addresses this by fitting all SNPs simultaneously within the genomic prediction framework, counteracting shrinkage and effect dilution, and then aggregating back-solved marker effects within LD-defined haplotype blocks (Shaffer et al., 2025). In addition, haplotypes are more useful for breeding because they represent stable, selectable chromosome segments rather than isolated SNPs, more sensitive to recombination. Block-level variance in haplotype effects provides a measure of the genomic importance of each region that can be more robust than single marker tests under these conditions. This approach was applied separately for SN and DTF across all six trials, enabling genomic regions acting on SN through phenology to be distinguished from those with direct effects on SN.

Trial-specific BLUPs (**ĝ**) for SN and DTF were extracted from their respective MET models and used to backsolve markers effects for 12 trial-specific analyses (six per trait). Marker effects𝛂̂)￼ for each trait in each trial were backsolved from the corresponding vector of genotype BLUPs (**ĝ**(Equation 9￼):

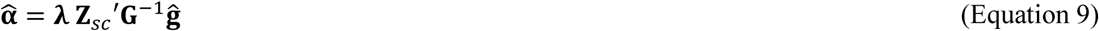

where 𝝀 is a scaling factor derived from allele frequencies across all loci, 𝐙_𝑠𝑐_′ is the transposed centered and scaled marker matrix and 𝐆^−1^ is the inverse of the GRM.

Haplotype blocks (haploblocks) were constructed independently for each of the eight chickpea chromosomes based on pairwise intra-chromosomal linkage disequilibrium (LD) between SNP markers (Shaffer et al., 2025). Blocks were defined using a flanking method, in which markers were iteratively added to both sides of a block when their r^2^ value with the nearest marker already included in the block met or exceeded the chromosome-specific LD threshold. If a marker failed to meet the threshold, the tolerance parameter determined how many subsequent markers were examined before the block was terminated. LD thresholds (r^2^ = 0.4 to 0.6) and tolerance values (1 to 2) were optimised independently for each chromosome by evaluating the number of blocks, proportion of single marker blocks, and mean and maximum markers per block, with parameters selected to balance robustness against imputation errors while maintaining detection power (Aldiss et al., 2025; Shaffer et al., 2025). Parameters for each chromosome are presented in Supplementary Table 8.

For each haploblock, the effect of each unique haplotype was estimated by summing the marker effects for all SNPs within the block. Local genomic estimated breeding values (localGEBVs) were then calculated for each genotype by assigning the haplotype effect corresponding to the genotype’s allelic configuration at each block. Block variance was calculated as the variance in unique haplotype effects across all genotypes, providing a measure of the genomic importance of each block for the trait of interest. The proportion of total block variance accounted for by the top 1% of haploblocks (n = 21) within each trial was calculated as the sum of block variances of the top 21 blocks divided by the sum of block variances across all 2109 blocks. To distinguish SN-specific blocks from those acting through phenology, the top 1% of haploblocks ranked by block variance for SN within each trial were classified by their overlap with the top 1% for DTF in the same trial. Blocks ranked in the top 1% for SN but not in the top 1% for DTF were classified as SN-specific in that trial.

To compare the genomic regions identified in the present study with previously reported loci, the physical positions of all top 1% SN, DTF and HSW blocks identified in this study were compared with the QTL reported by Jeffrey et al. (2024), which included trials conducted in environments partially overlapping with those used in the present study, although different traits were evaluated. The physical intervals of all top 1% SN, DTF and HSW blocks were compared with the published intervals of 14 QTL associated with yield, seed size, flowering time, maturity time, and final canopy closure to identify overlapping and unique genomic regions between the two studies. A genome-wide Circos plot was generated using the R package *circlize* to visualise the genomic distribution and overlap of QTL identified across both studies.

## 3. Results

### 3.1. A dual time of sowing strategy generated trials that experienced diverse environmental conditions

Sowing date manipulation combined with natural year to year variation produced six trials that spanned gradients of heat stress (0.0 to 13.2 days above 32°C), water availability (1 to 94 mm), and photosynthetic conditions (PTQ_VPD_ 0.96 to 1.86) during the critical period, providing a framework to evaluate genotypic response to diverse potential environmental conditions in one location (Table 2, Figure 2).

**Figure 2.**
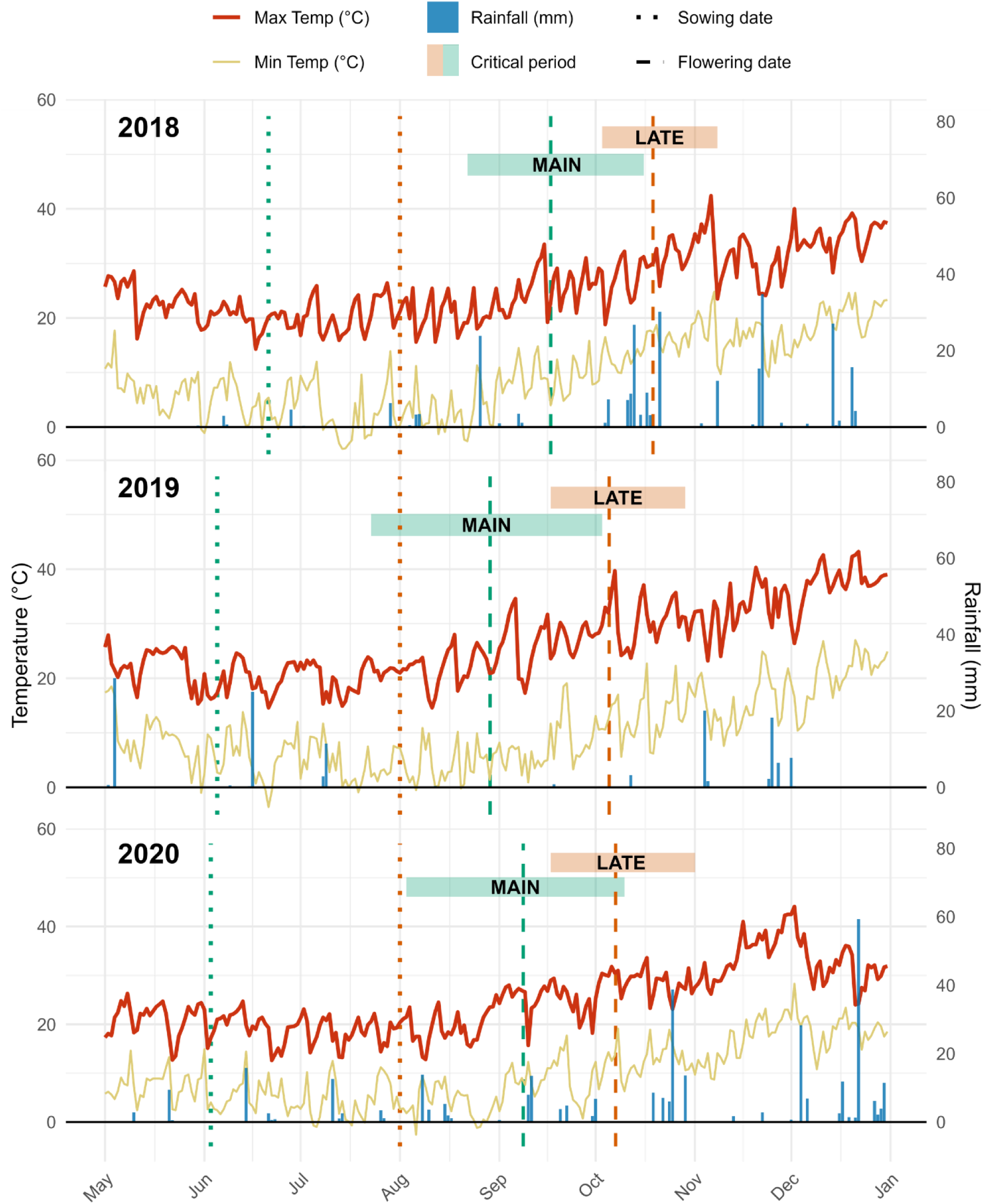
Daily maximum and minimum temperature (°C) and rainfall (mm) during six trials at Narrabri from 2018 to 2020. Green and orange represent MAIN and LATE trials respectively, and shaded boxes denote the mean timing and duration of the critical period. Dotted vertical lines indicate sowing date and dashed vertical lines indicate mean flowering date (expressed in days after sowing). Supplementary irrigation in 2019 is included as rainfall.

**Table 2.**
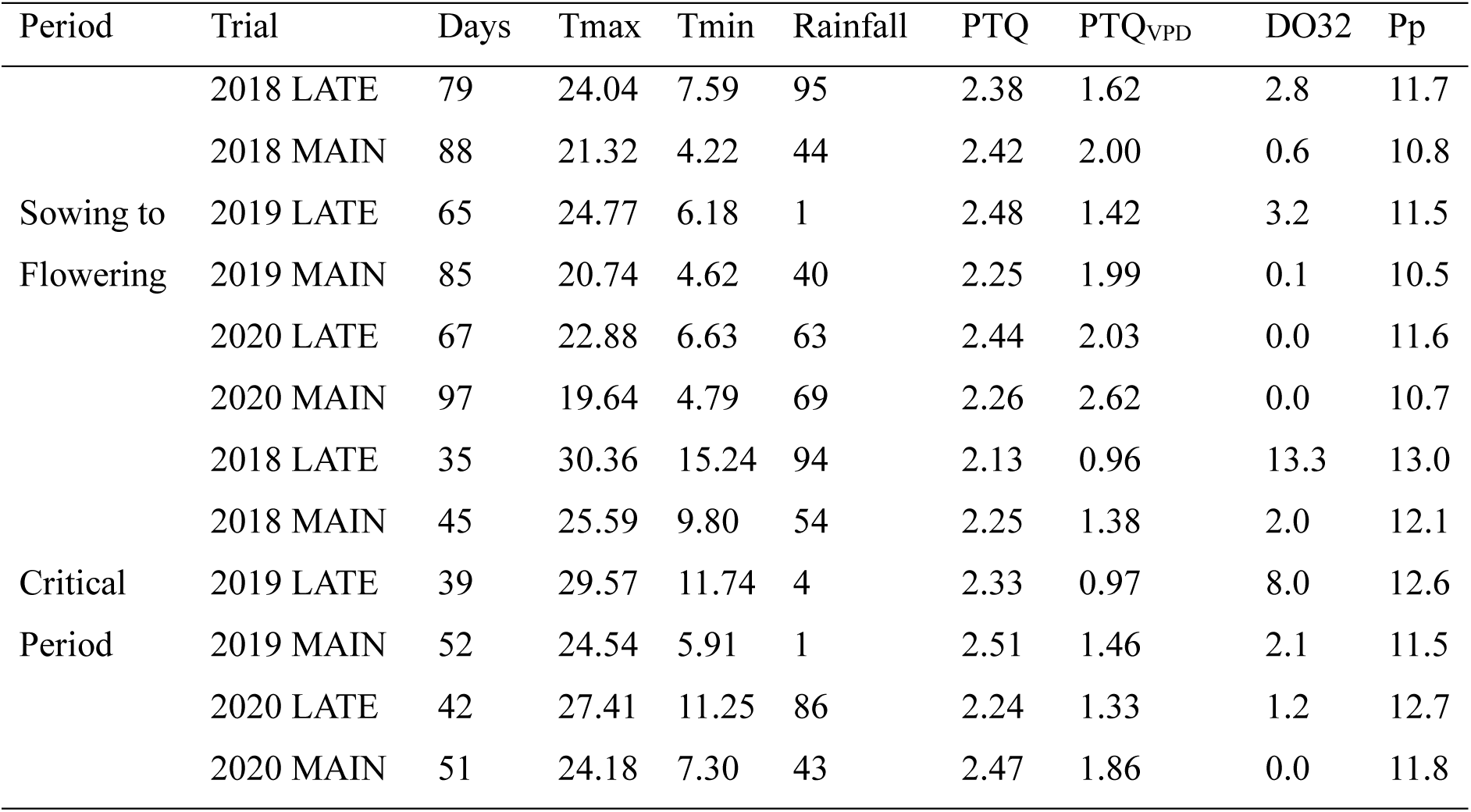
Summary of environmental characteristics during the sowing to flowering and critical period growth windows for six trials at Narrabri from 2018 to 2020. Trial means for the duration of each growth window in days (Days), mean daily maximum (Tmax; °C) and minimum (Tmin; °C) temperature, total rainfall (mm), photothermal quotient (PTQ; mol m⁻² °C⁻¹), photothermal quotient adjusted for vapour pressure deficit (PTQVPD; mol m⁻² °C⁻¹ kPa⁻¹), days above 32°C (DO32), and mean daylength (Photoperiod; h).

Overall, LATE trials had daily maximum temperatures 2.3 to 5.0°C higher during the critical period than MAIN trials. This temperature differential translated to more heat stress exposure, where LATE trials averaged 7.5 days above 32°C on average compared to 1.4 days in MAIN trials. The largest differential occurred in 2018, where the LATE trial genotypes experienced 13.3 days above 32°C on average relative to 2.0 days in the MAIN trial. Elevated temperatures in LATE trials resulted in faster thermal time accumulation which shortened the critical period in calendar days. The sowing to flowering window was compressed by 9 to 30 days in LATE relative to MAIN sowing across years, and the critical period by 6 to 10 days (Table 2, Figure 2). Despite this compression, LATE trials still accumulated more days above 32°C. Across all six trials, delayed sowing consistently elevated both temperature and atmospheric demand, reflected in lower PTQ_VPD_ values relative to MAIN trials in the same year.

Environmental covariates also had large seasonal variation. The 2018 season had the largest temperature differential between TOS, with adequate rainfall. The 2019 season had very low rainfall with MAIN and LATE trials receiving only 1 and 4 mm respectively during the critical period. The 2020 season had cooler growing temperatures where genotypes in the MAIN and LATE trials only experienced 0.0 and 1.2 days over 32°C during the critical period on average, respectively. This season also had the highest relative humidity and PTQ_VPD_ relative to 2018 and 2019.

### 3.2. Delayed sowing led to lower grain yield, despite genotype-dependent GEI

In each year, the LATE trial had a lower mean GY than the MAIN trial. The difference was -1.04 t ha^-1^ in 2018, -0.72 t ha^-1^ in 2019 and -0.64 t ha^-1^ in 2020. The mean GY was also lower each year from 2018 to 2020, in MAIN trials from 2.97 to 2.22 to 1.36 t ha^-1^ and in LATE trials from 1.94 to 1.50 to 0.72 t ha^-1^, respectively. The 2020 MAIN trial GY mean of 1.36 t ha^-1^ was lower than both the 2018 and 2019 LATE trials.

Substantial reranking of genotypes was observed for GY across the six trials in both commercial cultivars and the diverse germplasm panel (Figure 3a). Among 141 genotypes, 15 ranked above the trial mean in all six trials, including diverse accessions FLIP 94 62C (+0.35 t ha⁻¹ above trial mean on average), FLIP 09 333C (+0.32 t ha⁻¹), and FLIP 09 173C (+0.28 t ha⁻¹). Kimberley Large was the lowest ranked genotype in the entire panel (-0.64 t ha⁻¹ below trial mean on average), far lower performing than the next lowest genotype FLIP 09 356C (-0.30 t ha⁻¹). Almaz also yielded below the trial mean in five of six trials. In 2018, only Kimberley Large had a higher GY under delayed sowing (+0.29 t ha⁻¹; Figure 3b,c). In 2020, two genotypes FLIP 09 165C and FLIP 09 170C had a higher GY in the LATE sowing although the difference was small at +0.05 and +0.01 t ha⁻¹ respectively. No genotype showed higher GY from delayed sowing in 2019 (Figure 3a,b,c). Despite smaller absolute losses in 2020, the proportional reduction was greatest in that year due to the low MAIN trial GY (Figure 3b,c).

**Figure 3.**
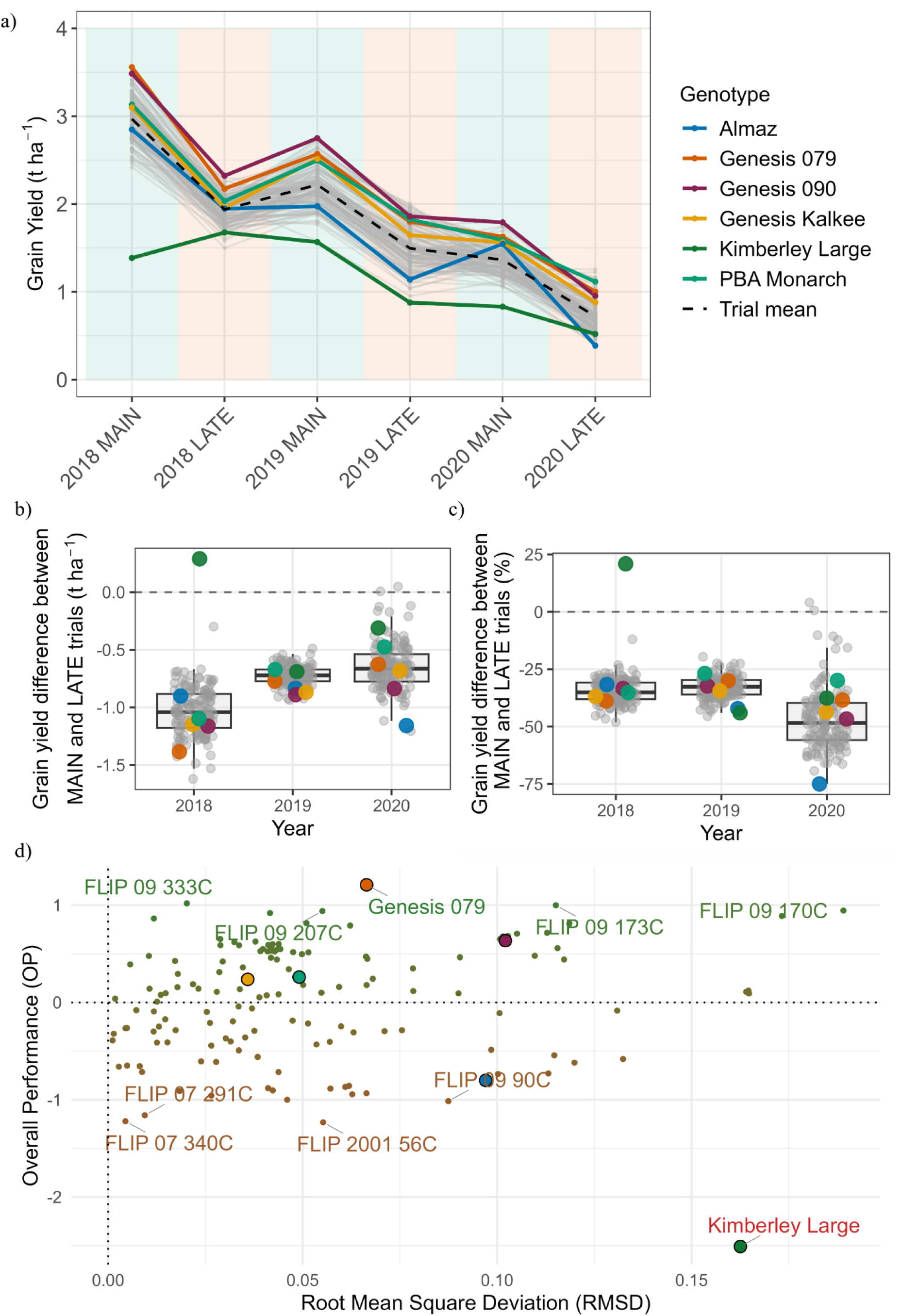
Genotype performance and GEI patterns for grain yield (GY; t ha⁻¹) in six trials at Narrabri from 2018 to 2020. a) Best Linear Unbiased Predictions (BLUPs) added to trial mean (μ) for GY plotted across trials. Green shading indicates MAIN trials and orange shading indicates LATE trials. Grey lines represent individual genotypes, coloured lines highlight six commercial Australian cultivars, and the black dashed line indicates the trial mean. b) Absolute difference and c) percentage difference in BLUPs between LATE and MAIN trials within the same year (LATE-MAIN). In panels b and c, boxplots show the distribution of genotypic differences, points indicate individual genotypes with commercial cultivars highlighted, and the dashed horizontal line indicates no difference in GY between time of sowing. d) Relationship between overall performance (OP) and root mean square deviation (RMSD) for BLUPs of GY across the six trials. Points are coloured on an OP gradient from green (highest) to red (lowest), filled circles highlight six commercial Australian cultivars, and the top and bottom five genotypes for OP are labelled.

Across all trials, 55% of genotypes achieved an above average overall performance (OP > 0). Genesis 079 was the highest performing genotype (OP = 1.21), followed by FLIP 09 333C and FLIP 09 173C (Figure 3). Among commercial cultivars, Genesis 079, Genesis 090, PBA Monarch, and Genesis Kalkee all had positive OP, while Kimberley Large had the lowest OP of all genotypes assessed. Some genotypes, such as FLIP 09 170C and FLIP 09 173C had a high RMSD indicating instability, despite being also among the top five genotypes for OP.

### 3.1. Grain yield is driven by seed number

#### 3.1.1. 100-seed weight is stable across environment while seed number captures GEI in grain yield

Genetic correlations between GY and SN were consistently positive and moderate to strong, ranging from 0.67 to 0.90 across trials. In contrast, genetic correlations between GY and HSW were negative and weaker, ranging from −0.47 to −0.09 (Table 3).

**Table 3.** Genetic correlations between grain yield (GY), and 100-seed weight (HSW) and seed number (SN), estimated from bivariate factor analytic models for six trials at Narrabri from 2018 to 2020.

| Trial | GY-SN | GY-HSW |
| --- | --- | --- |
| 2018 LATE | 0.73 | -0.21 |
| 2018 MAIN | 0.67 | -0.47 |
| 2019 LATE | 0.81 | -0.25 |
| 2019 MAIN | 0.79 | -0.34 |
| 2020 LATE | 0.90 | -0.09 |
| 2020 MAIN | 0.78 | -0.32 |

Genotype rankings for HSW were highly stable across environments where correlations ranged from 0.92 to 0.96 (Figure 4). This stability reflects the high additive variance of 72 to 99% of total genetic variance, and broad-sense heritability of 0.93 to 0.98 (Supplementary Table 3).

**Figure 4.**
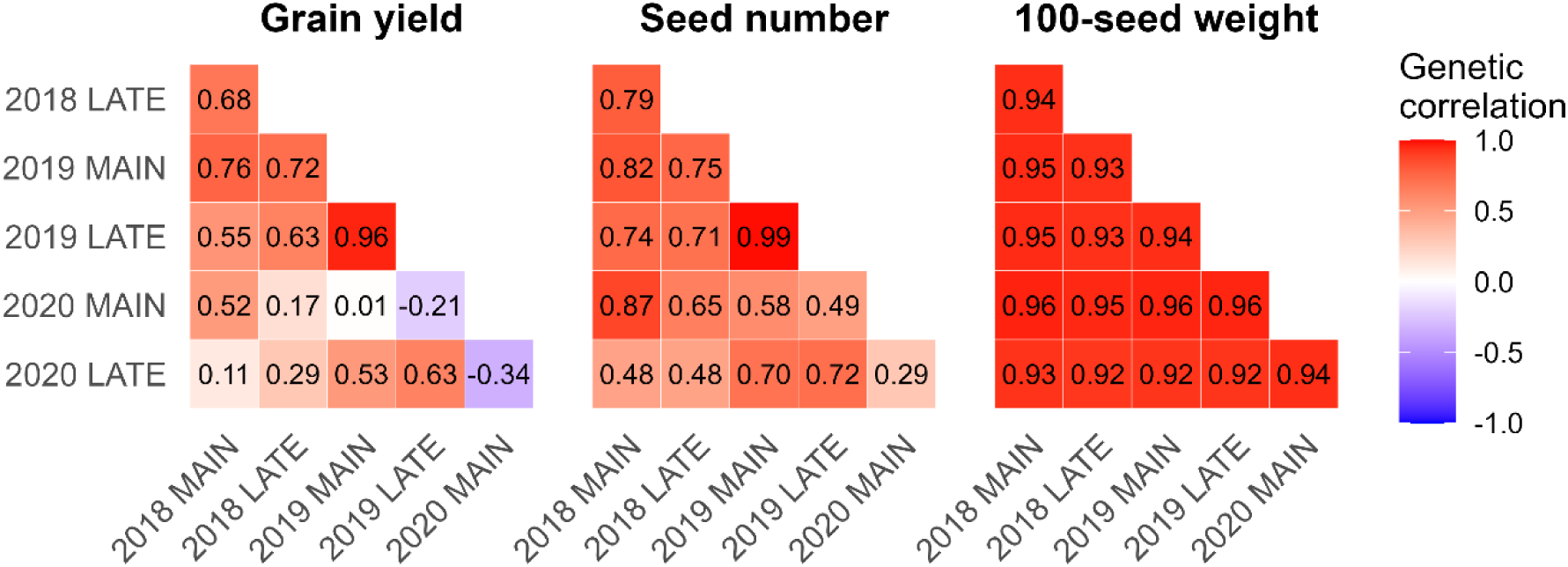
Genotype-by-environment correlations for grain yield (t ha^-1^), seed number (seeds m^-2^), and 100-seed weight (g) in six trials at Narrabri from 2018 to 2020. Values represent the genetic correlation of trait-specific effects between trial pairs, higher values indicate more stable genotype rankings across environments.

Greater GEI for SN was observed across trials, with genetic correlations from 0.29 to 0.99. The 2020 MAIN trial had the weakest correlations of 0.29 to 0.65 with other trials. Additive variance for SN ranged from 34 to 96% across trials (Supplementary Table 4). The greatest GEI for genotype rankings was observed for GY, ranging from −0.34 to 0.76, with the largest range observed in the 2020 MAIN trial of −0.34 to 0.52. The presence of GEI in both GY and SN, combined with their strong genetic correlation (Table 3), indicated that SN is a primary driver of environmental sensitivity in GY.

#### 3.1.2. Shorter phenological duration is associated with higher seed number

Delayed sowing accelerated phenological development in calendar days (Table 2), yet time to flowering, podding, and maturity was generally stable across trials when expressed in thermal time (Figure 5). Among all genotypes, the overall mean DTF was 1129°Cd, DTP was 1351°Cd, and DTM was 2037°Cd. Genetic variance for DTF was large in the 2018 LATE trial (Supplementary Table 5), with a range of 762°Cd (858 to 1620°Cd) compared to 243 to 360°Cd in all other trials (Figure 5c). Despite scale GEI, genotype rankings for DTF and DTP were stable across trials (Figure 5b, c), consistent with the high proportion of additive genetic variance (70 to 97% and 62 to 91% respectively) and FA1 models explaining 98.3% and 96.6% of additive genetic variance (Supplementary Table 5, Supplementary Table 6). Greater instability was observed in DTM, with crossover GEI particularly in the 2020 trials (Figure 5a), requiring an FA3 model to explain 86.5% of additive genetic variance. Despite podding occurring at a similar thermal time to other trials, the 2020 MAIN trial had the largest thermal time requirement for maturity, resulting in a long podding to maturity interval. Generally, commercial cultivars had a below average thermal time requirement to all phenological stages relative to the diversity panel, with the exception of Almaz which on average had a DTF, DTP and DTM above the trial mean, and Genesis Kalkee which also had above average DTF.

**Figure 5.**
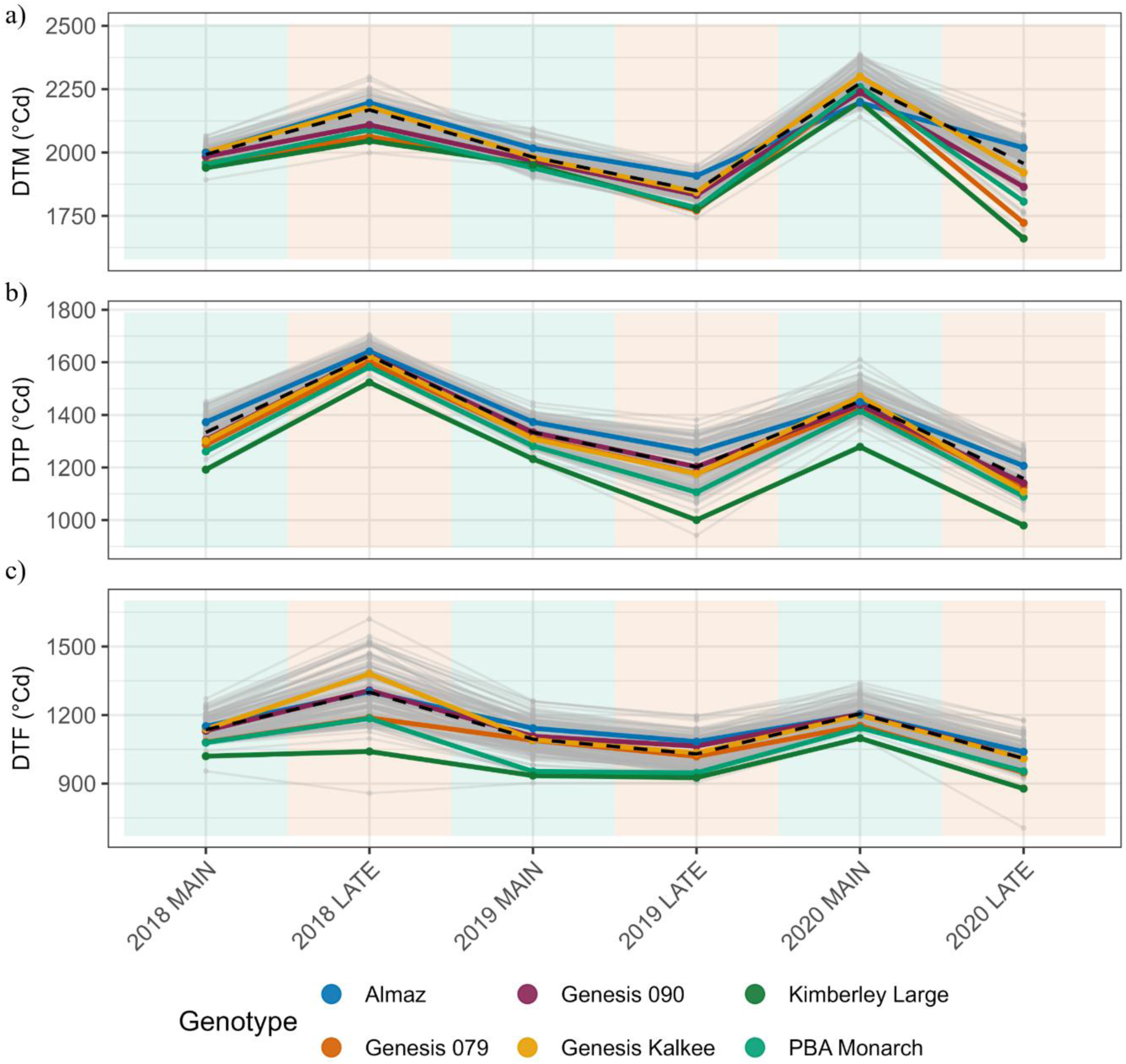
Best Linear Unbiased Predictions (BLUPs) added to trial mean (μ) of (a) thermal time to flowering (DTF), (b) thermal time to podding (DTP), and (c) thermal time to maturity (DTM) across six trials at Narrabri from 2018 to 2020. Proportion of additive variance explained by FA loadings were DTF (FA1, 98.3%) DTP (FA1, 96.6%) and DTM (FA3, 86.5%). Green shading indicates MAIN trials and orange shading indicates LATE trials, grey lines represent individual genotypes, coloured lines highlight commercial cultivars, and the black dashed line represents the trial mean.

A longer flowering window had a negative genetic correlation with SN in most trials. Except for the 2018 and 2020 MAIN trials, genetic correlations between SN and DTF, DTP, DTM ranged from -0.57 to -0.22. In the two MAIN trials with weak correlations, phenological variance was small, particularly for DTF (Supplementary Table 6, Supplementary Table 7).

### 3.2. Rare haplotypes for seed number in genebank accessions are absent from elite cultivars

A total of 2109 haploblocks were constructed across the chickpea genome (Figure 6). Block variance was calculated as the variance of haplotype effects within each haploblock and was used to rank blocks for each trait and trial. For SN, the 1% of blocks with the highest block variance (n = 21) explained 65.32% of the total block variance summed across all blocks in the 2018 LATE trial, 66.01% in 2018 MAIN, 61.48% in 2019 LATE, 63.08% in 2019 MAIN, 65.60% in 2020 LATE, and 68.62% in 2020 MAIN. Across the six trials, 43 unique haploblocks were identified in the top 1% by block variance for SN in at least one trial, and are presented in Supplementary Table 9. To distinguish between blocks that act on SN through pathways related to, and distinct from flowering, the top 1% of blocks for SN were classified within each trial by their overlap with the top 1% of blocks for DTF in the same trial.

**Figure 6.**
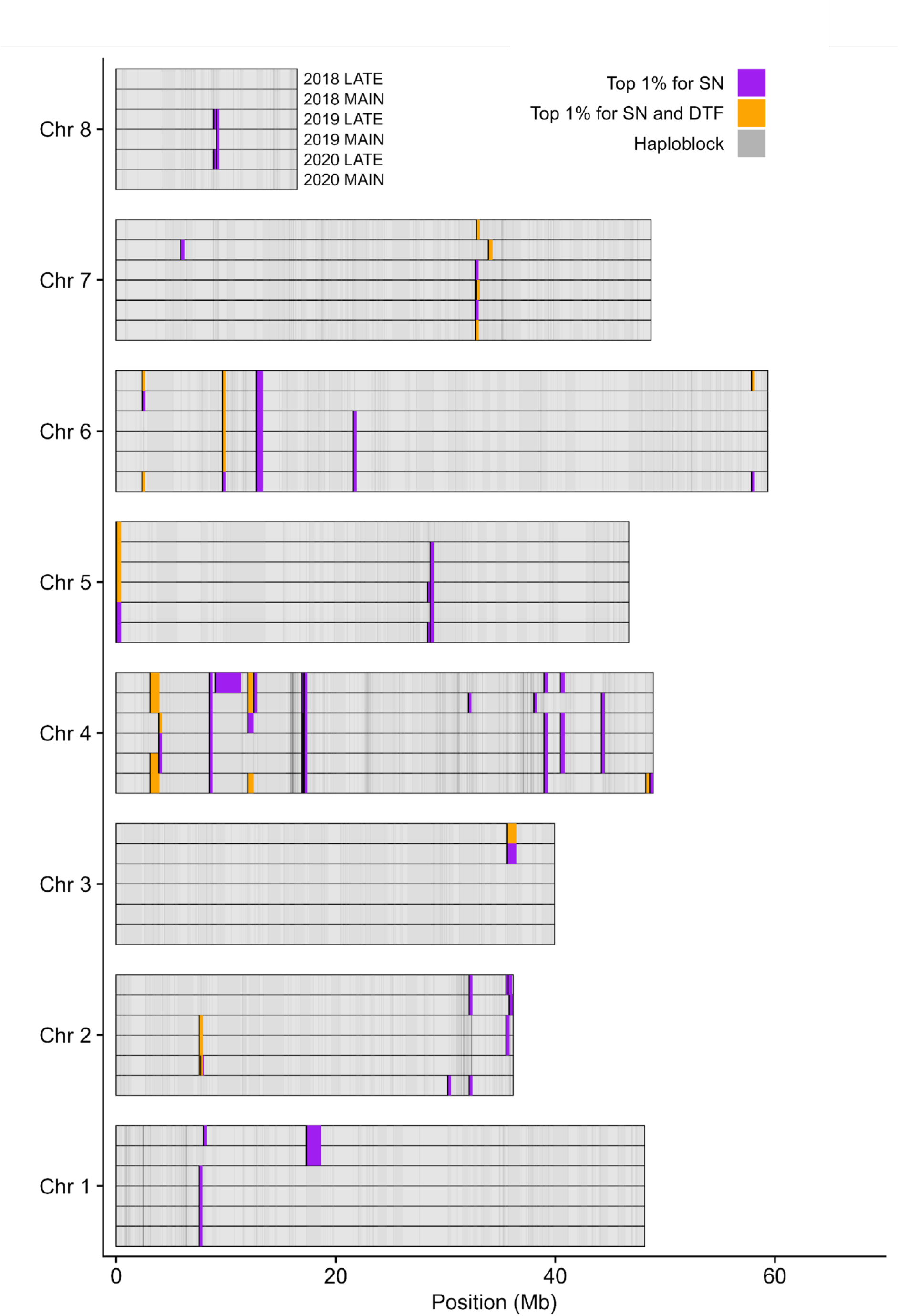
The top 1% of haploblocks with high block variance for seed number (SN) in six trials at Narrabri from 2018 to 2020, classified by overlap with the top 1% of haploblocks for thermal time to flowering (DTF) in the same trial. Block variance was calculated using Best Linear Unbiased Predictions (BLUPs) for each trait. Within each chromosome, rows correspond to the six trials. Purple segments show top 1% SN blocks that did not overlap with the top 1% for DTF in the same trial; orange segments show top 1% SN blocks that did overlap with the top 1% for DTF. Grey segments represent all other constructed haploblocks. Chromosomes are displayed along the y axis, with haploblock position and length drawn to scale on the x axis. Haploblocks smaller than 0.3 Mb have been artificially expanded to a minimum displayed width of 0.3 Mb for visibility.

Four blocks were identified in the top 1% for SN and not the top 1% for DTF in all six trials. Three were located on chromosome 4 (8.62 to 8.68 Mb, 16.97 to 17.19 Mb, and 17.25 to 17.26 Mb) and one on chromosome 6 (12.76 to 13.39 Mb). These SN-specific blocks represented the most robust candidates for targeted selection of SN independent of flowering time effects (Figure 7, Supplementary Figure 3).

**Figure 7.**
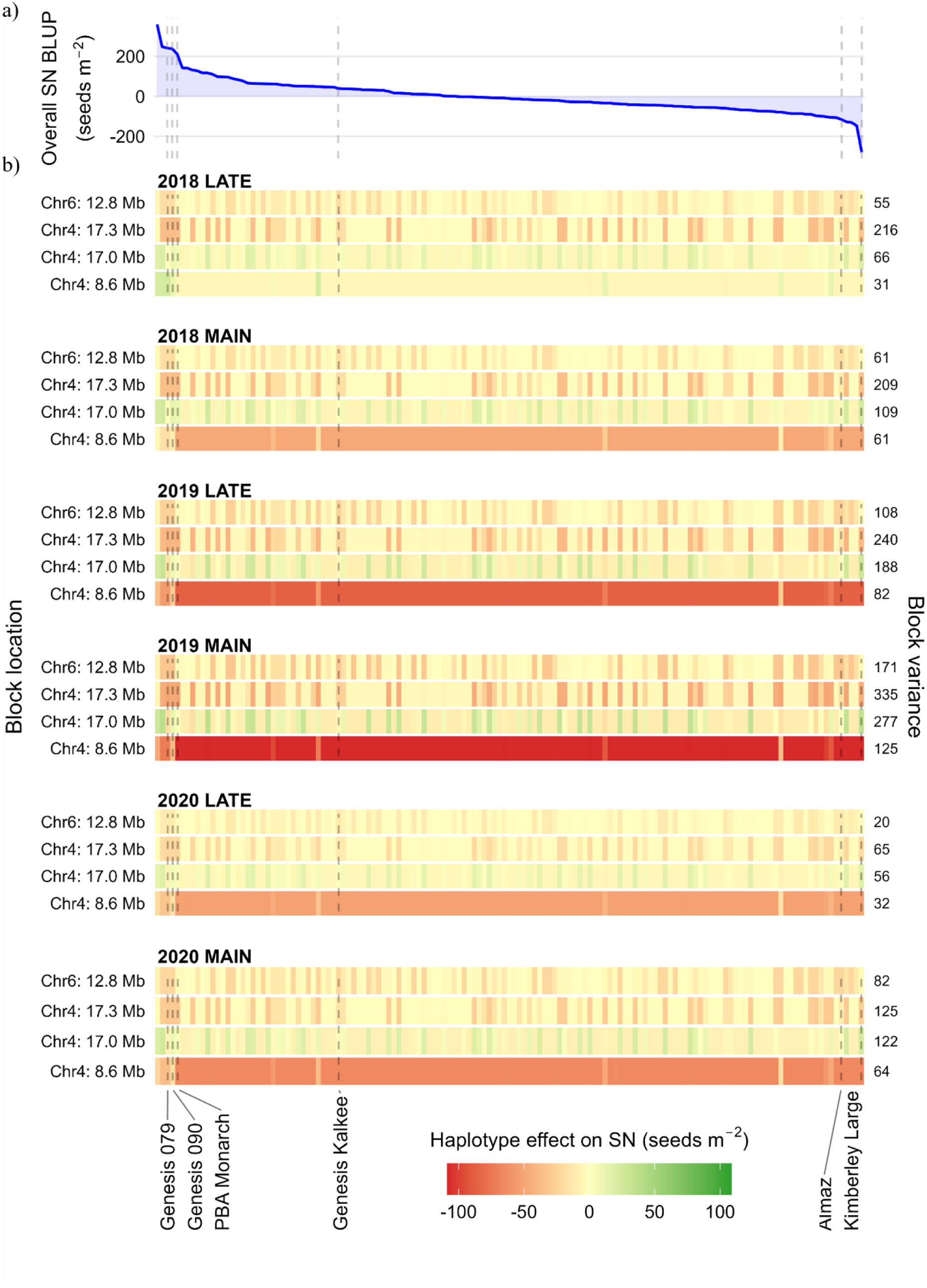
Haplotype effects on seed number (SN, seeds m^2^) at the four SN-specific haploblocks identified in all six trials at Narrabri from 2018 to 2020. (a) Best Linear Unbiased Predictions (BLUP) for SN across all six trials, with genotypes ordered from highest to lowest along the x axis. (b) Six heatmaps show haplotype effects (seeds m⁻²) at each of the four robust blocks, with one panel per trial. Each tile represents the haplotype effect carried by an individual genotype at the block, with the genotype order matching the BLUP ranking in the top panel. Green tiles indicate haplotypes with positive effects on SN, red tiles indicate negative effects, and yellow tiles indicate effects near zero. Colour intensity is on a common scale across all six trials. Right-hand y-axis values report block variance for each block within each trial. Vertical dashed lines indicate the position of six commercial Australian cultivars across all panels. Block coordinates are reported on the left y-axis.

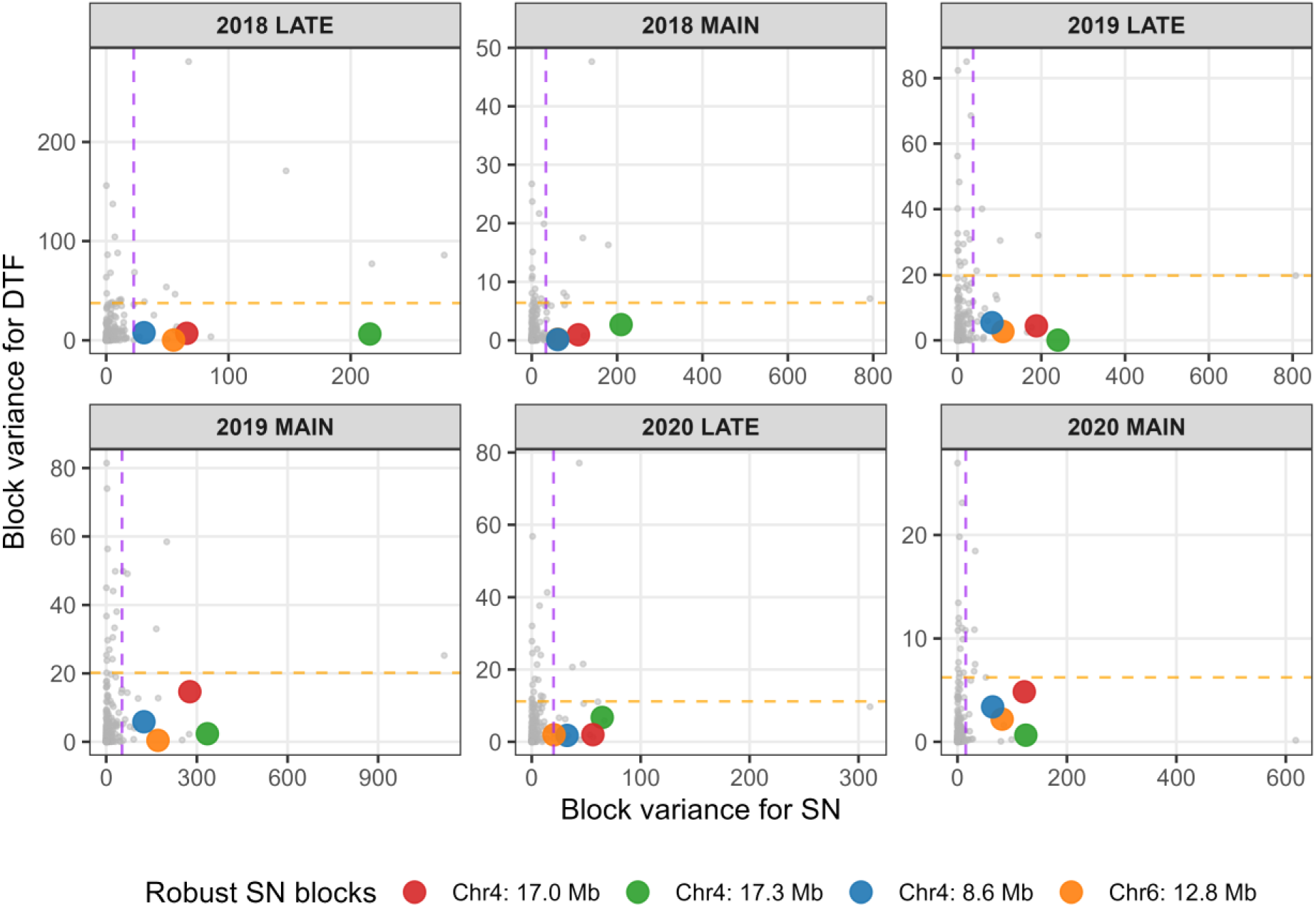

In addition, two top blocks were SN-specific in five of six trials, on chromosome 4 (38.97 to 39.34 Mb) and chromosome 5 (28.59 to 28.93 Mb). Five blocks were SN-specific in four of six trials, located on chromosome 1 (7.56 to 7.86 Mb), chromosome 4 (17.19 to 17.21 Mb, 40.44 to 40.88 Mb, and 44.30 to 44.38 Mb), and chromosome 6 (21.71 to 21.79 Mb). Six blocks consistently overlapped with the top 1% for DTF and were never classified as SN-specific. The most consistent of these was a block on chromosome 4 (3.10 to 3.95 Mb), identified in four trials, followed by blocks on chromosome 2 (7.72 to 7.73 Mb) and chromosome 6 (2.46 to 2.53 Mb), each identified in three trials. Three SN-specific blocks were identified only in the 2018 LATE trial, located on chromosome 1 (7.99 to 8.19 Mb), chromosome 2 (35.84 to 35.89 Mb), and chromosome 4 (9.01 to 11.38 Mb).

Comparison of the physical positions of top 1% blocks identified in the present study with the QTL intervals reported by Jeffrey et al. (2024) revealed eight overlapping genomic regions, including four QTL associated with seed size (*Qhsw_4.1*, *Qhsw_4.2*, *Qhsw_4.3*, *Qhsw_4.4*), two with maturity time (*Qmat_4.1*, *Qmat_8.2*), one with flowering time (*Qflo_6.2*), and one with canopy closure (*Qcanopy_1.2*) (Supplementary Figure 4).

Haplotype effects at the four robust SN-specific blocks revealed differences in selection potential across the six trials (Figure 7). Block variance was highest at the chromosome 4 blocks at 17.3 Mb (range 65 to 335) and 17.0 Mb (range 56 to 277), and lower at the blocks at 8.6 Mb (31 to 125) and chromosome 6 at 12.8 Mb (20 to 171). The chromosome 4 block at 17.0 Mb was the strongest candidate for targeted selection on SN. This block contained 30 unique haplotypes across the panel. The same haplotype was identified as the highest effect haplotype in all six trials, with consistently positive effects ranging from

+17.9 to +37.6 seeds m⁻². However, this most superior haplotype was carried by only two genotypes, FLIP 94 62C and FLIP 09 404C. The chromosome 4 block at 17.3 Mb contained 23 unique haplotypes and similarly showed consistent direction of effect across all six trials, with the same haplotype identified as the highest effect in every trial. The haplotype effects ranged from -49.9 to -0.6 seeds m⁻² across the panel, with the most superior haplotype carried by 79 genotypes (56%). The chromosome 6 block at 12.8 Mb showed the smallest haplotype effects of the four blocks. The haplotype with most positive effect across the six trials was near neutral (range -0.12 to +0.04 seeds m⁻²) and was carried by 40 genotypes (28%), while remaining genotypes carried haplotypes with an effect ranging down to -31.9 seeds m⁻². In contrast, the chromosome 4 block at 8.6 Mb showed GEI across trials. At this block, 123 genotypes (87.2%) carried the same haplotype, the remaining 18 genotypes had 13 different haplotypes. The haplotype with the highest effect differed between trials. In the 2018 LATE trial, the highest effect haplotype was rare in the panel, carried by only FLIP 94 62C and Genesis 079, and had a positive effect on SN (+24.0 seeds m⁻²). In all five other trials, this rare haplotype was not the most positive effect haplotype, and all haplotypes at the block had a negative effect on SN. Among the genotypes evaluated in this study, FLIP 94 62C was the only genotype to carry the rare superior haplotypes at both blocks (chromosome 4 at 17.0 Mb and 8.6 Mb).

## 4. Discussion

Matching cultivar to management and environment is a fundamental goal of plant breeding to achieve high and stable yield. In this study, delayed sowing reduced GY by up to 1.04 t ha⁻¹ and GEI for GY was driven largely by variation in SN and not HSW. Furthermore, accelerated flowering was a key component of adaptation among commercial cultivars, and genomic targets for SN improvement were identified on chromosomes 4 and 6. The high performance of several commercial cultivars across both times of sowing (TOS) and years suggests that long term selection across Australia’s broad production landscape has indirectly conferred resilience to the environments present in this analysis.

### 4.1. Strengths and limitations of delayed sowing as a screening tool

Delayed sowing in this study exposed plants to combinations of heat stress, rainfall, and elevated atmospheric demand. This combined stress is characteristic of field conditions, rather than controlled experiments that isolate single stressors (Kaushal et al., 2013; Krishnamurthy et al., 2011; Lake et al., 2025). Compared with controlled environment studies, delayed sowing better represents commercial production where stressors interact dynamically (Tardieu & Tuberosa, 2010). It is also a practical means to screen large germplasm collections in grower relevant conditions. Although kabuli chickpea is typically not grown in Narrabri, genetic resources identified here are relevant for introgression into elite desi backgrounds, and for expansion of kabuli varieties into this TPE. The 2018 LATE trial had the largest heat stress exposure on average among these trials, with 13.3 days above 32°C during the critical period. This occurred despite the smallest sowing date offset of 41 days, compared with 57 and 59 days in 2019 and 2020 (Table 1, Table 2). In contrast, the 2020 LATE trial recorded only 1.2 days above 32°C. Delayed sowing reliably produces the combination of stressors characteristic of commercial production, but the severity of heat stress is not guaranteed and varies considerably across seasons, as demonstrated by the contrast between the 2018 and 2020 LATE trials in this study.

The critical period window applied here was derived from a limited number of commercial cultivars (Lake & Sadras, 2014), and since the critical period was defined using this fixed window, diverse accessions may have genotype-specific windows that differ from those assumed here, potentially introducing misclassification of stress exposure for some genotypes. Additionally, the critical period was defined relative to the flowering date of each plot. Since flowering dates clustered within a small window and seasonal conditions changed slowly relative to the spread in flowering time, the range of environmental covariate values across plots within a trial remained narrow. When phenological variation among germplasm is limited, relationships between genotype performance and environmental covariates relative to phenology are difficult to detect. Future analyses may explicitly include environmental covariates in the statistical model to partition GEI (Costa-Neto et al., 2021; Jarquín et al., 2014).

The six trials in this study spanned a range of environmental conditions, representative of the Northern Grain Region TPE (Chauhan et al., 2017; Dreccer et al., 2018). Regardless, the single location in this study limits extrapolation of genotype rankings and haploblock effects across a broader TPE. Further, the trial density (three years, two times of sowing) was insufficient to formally characterise the TPE or distinguish consistent environment types from year to year variation (Supplementary Figure 2). Future experiments may consider local historical data and incorporate multiple years and locations to increase the likelihood of capturing agronomically relevant stress exposure (Chauhan et al., 2017; Lake et al., 2025), or consider irrigation to limit the confounding effect of soil moisture on heat stress response.

### 4.2. Genotype reranking across environments was driven by seed number, while seed weight was stable

In this study, HSW had high genetic correlations across all trials (Figure 4), high additive genetic variance and broad-sense heritability (Supplementary Table 3), consistent with previous analysis of the dataset (Jeffrey et al., 2025). Fewer environments are therefore required to phenotype and assess merit for this trait. However, HSW does not reflect other seed quality traits including colour, shape, size uniformity, and defects (e.g. shrivelled grain) that are of critical economic importance and have unique responses to stress (Wood & Scott, 2021).

In contrast, GY had significant reranking across environments (Figure 4). It is important to note that SN was calculated from GY and HSW rather than measured directly, which limits biological inference on whether reductions in SN were driven by fewer pods through flower or pod abortion, or fewer seeds per pod through seed abortion. Seed number had strong positive genetic correlations with GY across all trials while HSW had weak to moderate negative correlations. Despite HSW rankings being highly stable across environments, the GEI in GY was captured primarily by variation in SN rather than seed weight. While the mean HSW was higher in MAIN trials than LATE trials, the difference was small at 2.70 to 5.07 g, relative to the observed GY reductions of 0.64 to 1.04 t ha⁻¹. This indicates that GY losses under these environments were driven primarily by reductions in SN rather than seed weight, consistent with previous findings in chickpea and other pulse crops (Jeffrey et al., 2025; Pushpavalli et al., 2015; Sadras et al., 2015; Sandana & Calderini, 2012). The stability of HSW likely reflects physiological adaptive mechanisms including assimilate partitioning to maintain viable seed weight, and flower or pod abortion under stress (Fang et al., 2010; Leport et al., 2006; Nakano et al., 1998).

Genetic correlations for GY were lower than SN genetic correlations in some trial pairs, so additional sources of GEI not captured by SN alone also contributed to GY variation. These may include plot level effects such as plant establishment, lodging or harvest losses.

Four of six commercial cultivars had an above average OP across the six trials (Figure 3). Kimberley Large, a cultivar adapted to the irrigated production regions of the Kimberley area in Western Australia and Central Queensland (Siddique & Regan, 2005), had the lowest OP overall. However, it was the only genotype to show a GY advantage under delayed sowing in 2018, potentially reflecting adaptation to warmer growing conditions in its TPE. Almaz, a cultivar developed for winter sowing with mild spring conditions (Siddique et al., 2007), underperformed under delayed sowing with a 75% GY reduction in 2020 (Figure 4), consistent with poor adaptation to the hot spring conditions of northern NSW. These outcomes illustrate the importance of maintaining well characterised TPE and regional variety testing to avoid misalignment between cultivar adaptation and production system, supported by the distinct genomic backgrounds of Kimberley Large and Almaz relative to the other commercial cultivars evaluated in this study (Figure 1).

Commercial cultivars had faster flowering than average among the panel (Figure 5) which suggests that shorter flowering duration has been an important component of their adaptation strategy. Small variation in DTF among commercial cultivars relative to the broader germplasm panel suggests that while current cultivars are well adapted to existing conditions, they may lack the genetic diversity needed to respond to increasingly variable climates (Figure 5).

Warmer temperatures due to delayed sowing accelerated phenological development in calendar days (Table 2, Figure 2), consistent with established temperature responses in chickpea (Soltani & Sinclair, 2011). Although the thermal time model assumed linear temperature responses and did not incorporate photoperiod, soil water, or radiation effects (Chauhan et al., 2019; Gimenez et al., 2025; Li et al., 2022), the narrow photoperiod range across trials (10.5 to 11.7 hours from sowing to flowering) and consistency of thermal time requirements for developmental stages across environments (Figure 5) indicates adequate performance for comparative purposes. Genetic rankings for DTF and DTP remained highly stable across trials, indicating that selection in any single trial would effectively predict rankings across the panel.

Greater complexity emerged at maturity, which required an FA3 model to explain 86.5% of additive variance (Supplementary Table 7). The podding to maturity interval was more susceptible to non-additive and trial specific effects, which may reflect environmental signals such as water status, canopy senescence dynamics, and source-sink relationships that have interactions specific to each genotype. Earlier pod set has been shown to increase harvest index in chickpea, particularly under terminal drought environments where a longer pod setting phase compensates for reduced biomass (Gimenez et al., 2025). Across most trials, an extended phenology was associated with lower SN (Table 4), consistent with shorter duration phenology providing escape from terminal stress. The weak correlations between SN and phenology in the 2018 and 2020 MAIN trials likely reflect the limited variance in phenology among genotypes (Supplementary Table 5, Supplementary Table 6).

**Table 4.** Genetic correlations between seed number (SN) and thermal time to flowering (DTF), podding (DTP), and maturity (DTM), estimated from bivariate factor analytic models for six trials at Narrabri from 2018 to 2020.

| Trial | SN-DTF | SN-DTP | SN-DTM |
| --- | --- | --- | --- |
| 2018 LATE | -0.29 | -0.24 | -0.37 |
| 2018 MAIN | 0.11 | 0.09 | 0.03 |
| 2019 LATE | -0.44 | -0.29 | -0.55 |
| 2019 MAIN | -0.35 | -0.36 | -0.57 |
| 2020 LATE | -0.22 | -0.26 | -0.36 |
| 2020 MAIN | 0.08 | 0.14 | -0.35 |

### 4.3. Broadening the genetic base through introgression of rare haplotypes from genebank accessions

Although kabuli chickpea commands a price premium over desi types due to its larger seed size and suitability for whole grain markets, kabuli constitutes only ∼15% of Australian chickpea plantings due to limited export markets and production constraints (Hobson et al., 2016; Subedi et al., 2024). Identification of genetic resources in kabuli germplasm presents opportunities for both variety development within the kabuli gene pool to improve adaptation, and for introgression of novel heat adaptive haplotype blocks into elite desi backgrounds.

This study extends the genomic findings of Jeffrey et al. (2024), who identified 14 QTL associated with yield, seed size, flowering time, maturity time, and final canopy closure using GWAS on germplasm that overlaps with the panel evaluated here. The localGEBV approach identified a larger number of genomic regions, with 105 unique top 1% blocks across all environments, including 43 associated with SN, 40 associated with DTF, and 22 associated with HSW. This greater resolution reflects a fundamental difference in methodology. While GWAS tests markers individually, the localGEBV approach fits all SNPs simultaneously, concentrating statistical power within LD-defined haplotype blocks and improving detection sensitivity, particularly in panels of moderate size (Shaffer et al., 2025). The overlapping regions identified between both Jeffrey et al. (2024) and this study suggest shared genetic control of SN, developmental timing, and other agronomic traits in chickpea, and provide independent validation of these genomic regions across different analytical frameworks. Six previously reported QTL were not detected in the present study, which may reflect differences in germplasm and environments evaluated, traits assessed, or the distinct statistical power profiles of GWAS versus localGEBV approaches.

A challenge with field trials is to disentangle the direct effects of yield components on grain yield, from the indirect effects of environment and phenology (Sadras et al., 2015). Classification of SN haploblocks by their overlap with DTF haploblocks allowed identification of high variance regions that act on SN through pathways potentially independent of phenology. Flowering was used as the phenology trait for haplotype classification because it anchored the critical period (Lake & Sadras, 2014), had high broad-sense heritability and low GEI (Figure 5, Supplementary Table 5). The 1% threshold used here was for exploratory purposes however for breeding applications a threshold should be determined by what is considered a meaningful agronomic outcome.

The greatest concentration of high variance blocks for SN was on chromosome 4, consistent with reports of a QTL cluster for yield and drought adaptive traits on this chromosome (Barmukh et al., 2022; Garg et al., 2025; Nguyen et al., 2022; Varshney et al., 2014). However, the blocks identified here are different from the QTL hotspot region defined at 13.2 to 13.5 Mb (Kale et al., 2015), which suggests these may represent independent genomic regions associated with SN performance under the environments assessed. Notably, none of the four most stable and robust blocks identified in the present study overlapped with the QTL reported by Jeffrey et al. (2024), further highlighting the capacity of the localGEBV framework to resolve novel genomic regions that complement those identified by single-marker approaches.

Six independent analyses each identified four haploblocks in the top 1% of high variance for SN, which provides confidence that they are not artefacts of a single trial. The two blocks on chromosome 4 at 17.0 and 17.3 Mb were spatially adjacent (68 kb apart), and while the LD criterion treated them as statistically separate based on direct correlation between SNPs, both blocks may be in LD with the same unobserved causal variant. Higher marker density or fine mapping may resolve whether these represent one or two distinct genes.

Examination of haplotype effects at the four blocks revealed that high block variance does not always translate to value for breeding. In two blocks, the most superior haplotype was already present in 56% (chromosome 4 17.3 Mb) and 28% (chromosome 6 12.8 Mb) of genotypes. In the other two blocks (chromosome 4 8.6 Mb and 17.0 Mb), the most superior haplotype was present in only two genotypes, offering an opportunity to introgress rare allelic combinations for genetic gain.

A particularly important finding was the heat-specific response at the chromosome 4 block at 8.6 Mb. The rare haplotype carried by FLIP 94 62C and Genesis 079 conferred a positive effect on SN exclusively in the 2018 LATE trial, which was the trial with the greatest heat stress exposure, recording 13.3 days above 32°C during the critical period (Table 2). In all five other trials, including those with minimal heat stress, this haplotype had a negative effect on SN, and a different haplotype was most superior. This crossover interaction identifies chromosome 4 at 8.6 Mb as a candidate region for heat-specific SN improvement, distinct from regions conferring broad adaptation. It also demonstrates that selection for heat tolerance at this block requires targeted screening under high-temperature conditions, as multi-environment evaluation across benign environments would mask or select against the beneficial allele. This block represents a priority for further investigation and validation across additional heat stress environments.

### 4.4. Conclusion

In this study, delayed sowing reduced grain yield in kabuli chickpea primarily through reductions in seed number rather than seed weight. Accelerated phenology has supported adaptation in elite Australian cultivars relative to diverse germplasm. Rare superior haplotypes for seed number identified in genebank accessions provide an opportunity to improve and broaden the genetic base of Australian kabuli chickpea.

## 5. CRediT authorship contribution statement

C.B.J., SVH, LTH, RT, MRS: conceptualisation. LZ: data curation. C.B.J., SVH, VP, AK: formal analysis. RT, MRS; funding acquisition. CJ, LZ, RT: investigation. SVH, LTH, KC, AK, MRS; supervision. C.B.J.; writing - original draft, visualisation. C.B.J., SVH, LZ, CJ, VP, AK, RT, LTH, KC, MRS; writing - review and editing.

## 8. Acknowledgements

C.B.J. is an affiliate of the ARC Training Centre in Predictive Breeding for Agricultural Futures (IC230100016). The authors would like to thank Professor Graeme Hammer for his valuable input.

## 9. Data availability

Data that support the findings of this study are available on request.

## 10. Appendix A. Supplementary Material

**Supplementary Table 1.**
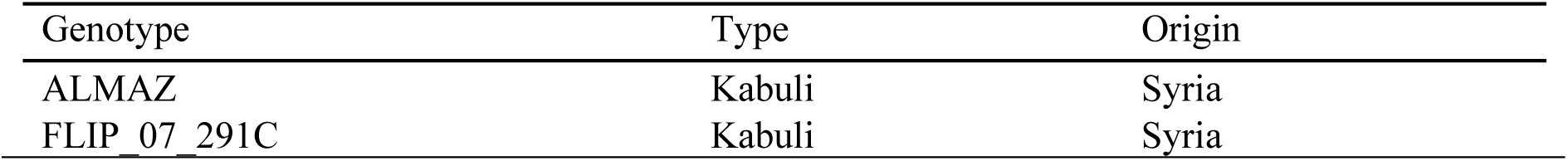
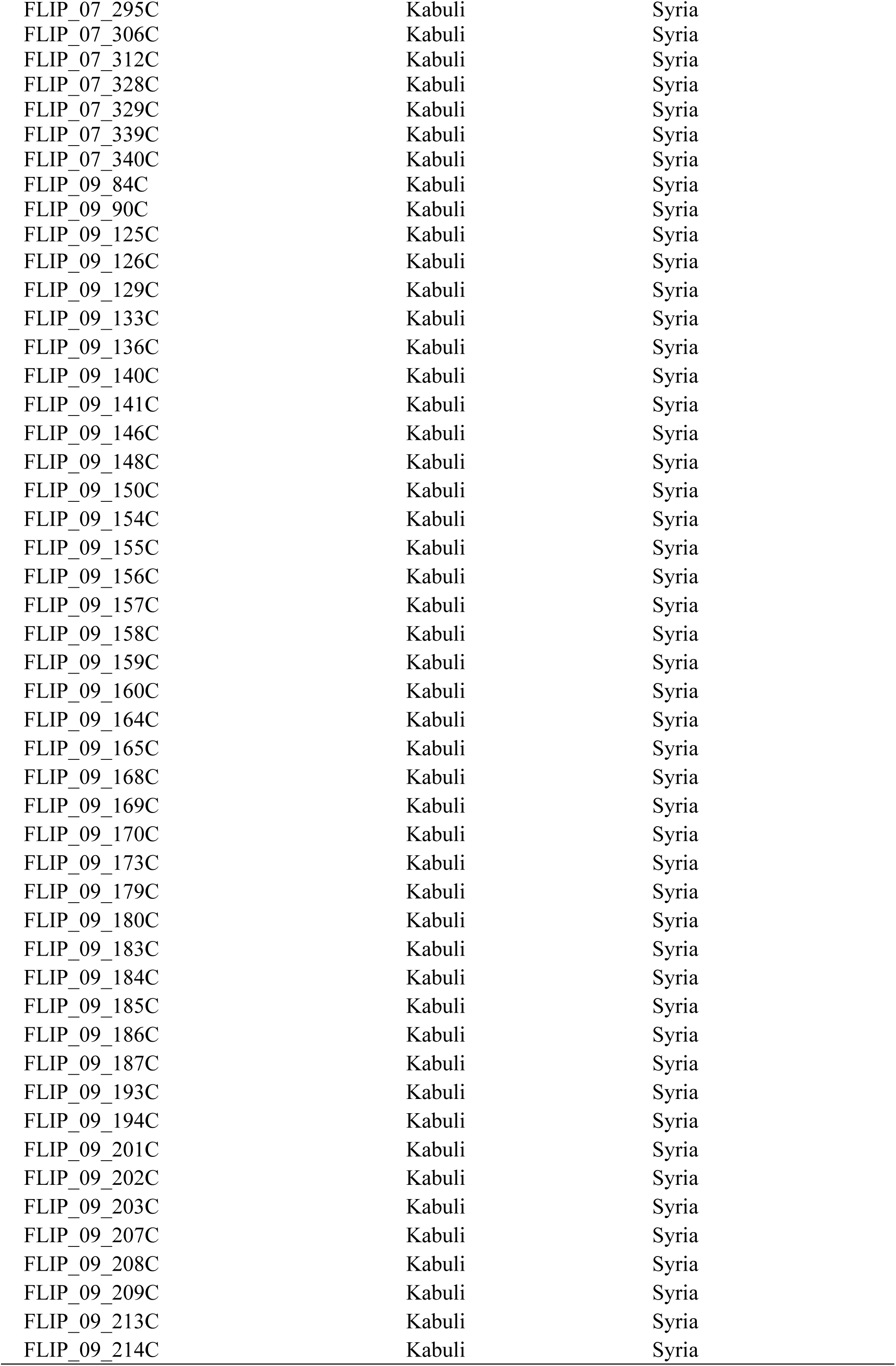
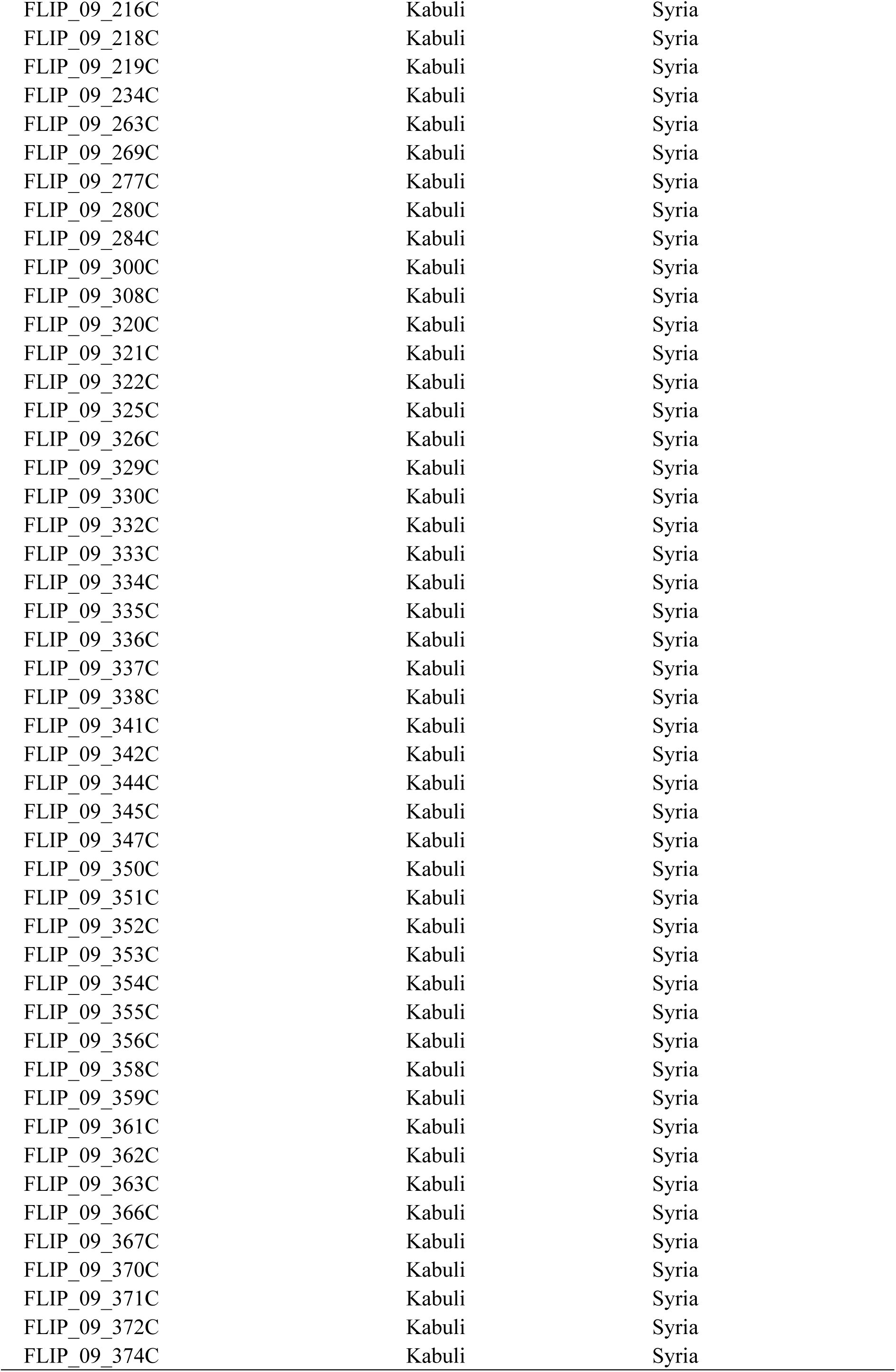
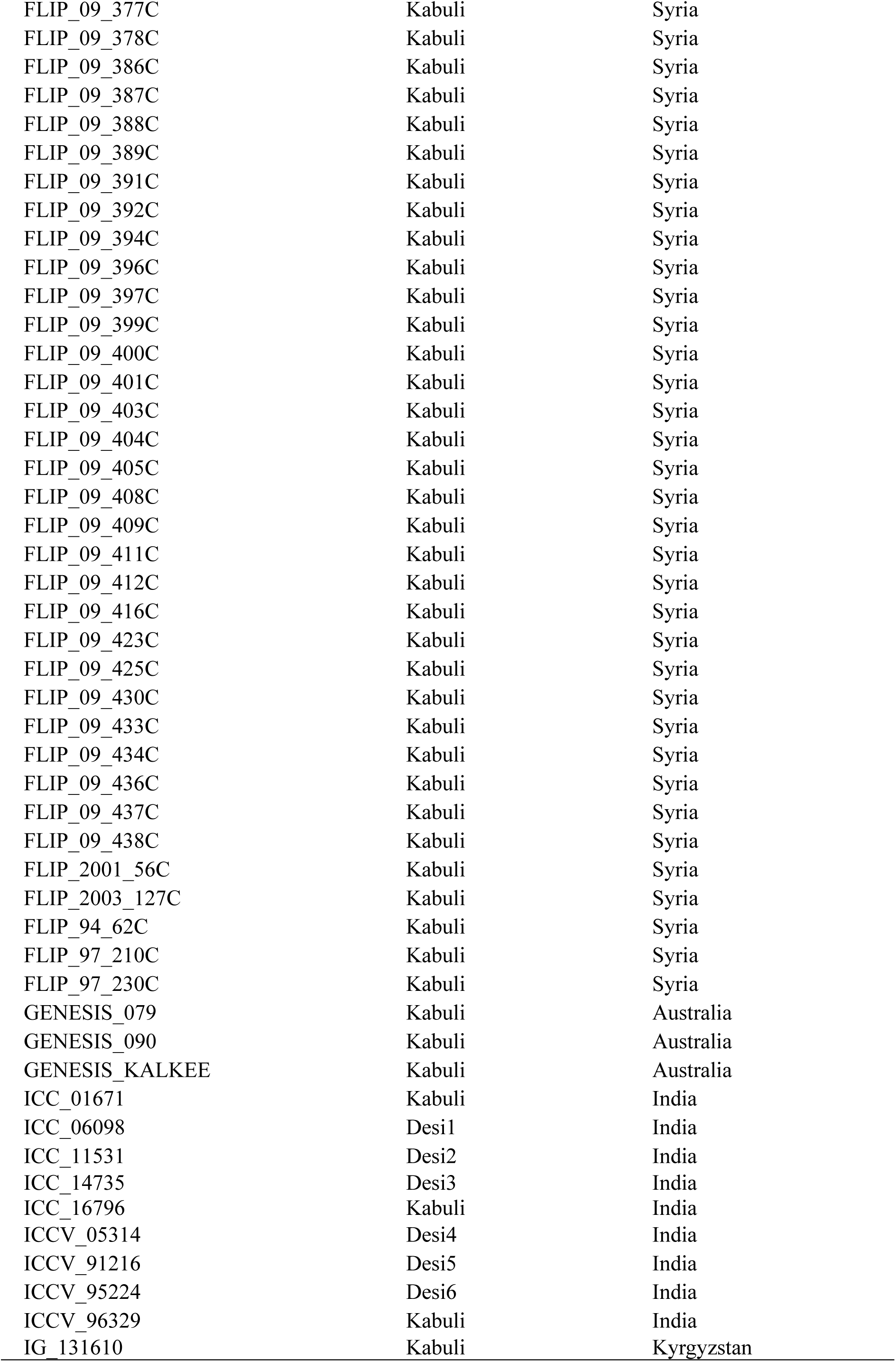
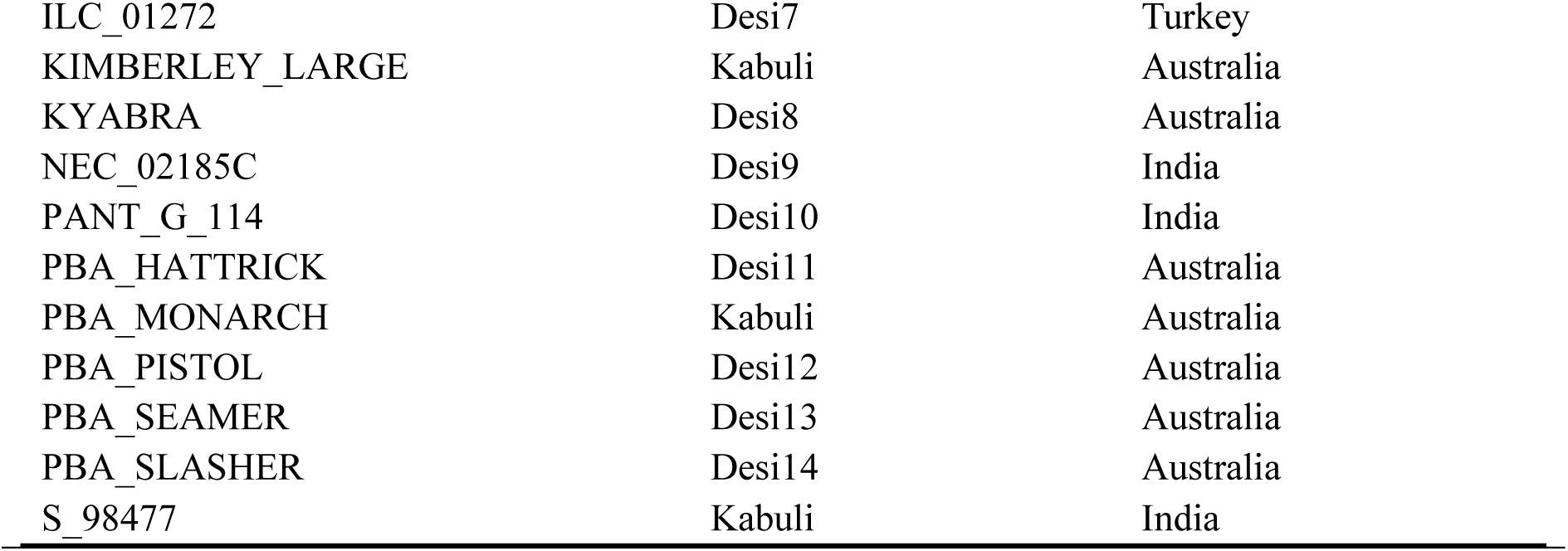
List of 157 unique genotypes, their type and origin, included in six trials at Narrabri from 2018 to 2020

| Genotype | Type | Origin |
| --- | --- | --- |
| ALMAZ | Kabuli | Syria |
| FLIP_07_291C | Kabuli | Syria |
| FLIP_07_295C | Kabuli | Syria |
| FLIP_07_306C | Kabuli | Syria |
| FLIP_07_312C | Kabuli | Syria |
| FLIP_07_328C | Kabuli | Syria |
| FLIP_07_329C | Kabuli | Syria |
| FLIP_07_339C | Kabuli | Syria |
| FLIP_07_340C | Kabuli | Syria |
| FLIP_09_84C | Kabuli | Syria |
| FLIP_09_90C | Kabuli | Syria |
| FLIP_09_125C | Kabuli | Syria |
| FLIP_09_126C | Kabuli | Syria |
| FLIP_09_129C | Kabuli | Syria |
| FLIP_09_133C | Kabuli | Syria |
| FLIP_09_136C | Kabuli | Syria |
| FLIP_09_140C | Kabuli | Syria |
| FLIP_09_141C | Kabuli | Syria |
| FLIP_09_146C | Kabuli | Syria |
| FLIP_09_148C | Kabuli | Syria |
| FLIP_09_150C | Kabuli | Syria |
| FLIP_09_154C | Kabuli | Syria |
| FLIP_09_155C | Kabuli | Syria |
| FLIP_09_156C | Kabuli | Syria |
| FLIP_09_157C | Kabuli | Syria |
| FLIP_09_158C | Kabuli | Syria |
| FLIP_09_159C | Kabuli | Syria |
| FLIP_09_160C | Kabuli | Syria |
| FLIP_09_164C | Kabuli | Syria |
| FLIP_09_165C | Kabuli | Syria |
| FLIP_09_168C | Kabuli | Syria |
| FLIP_09_169C | Kabuli | Syria |
| FLIP_09_170C | Kabuli | Syria |
| FLIP_09_173C | Kabuli | Syria |
| FLIP_09_179C | Kabuli | Syria |
| FLIP_09_180C | Kabuli | Syria |
| FLIP_09_183C | Kabuli | Syria |
| FLIP_09_184C | Kabuli | Syria |
| FLIP_09_185C | Kabuli | Syria |
| FLIP_09_186C | Kabuli | Syria |
| FLIP_09_187C | Kabuli | Syria |
| FLIP_09_193C | Kabuli | Syria |
| FLIP_09_194C | Kabuli | Syria |
| FLIP_09_201C | Kabuli | Syria |
| FLIP_09_202C | Kabuli | Syria |
| FLIP_09_203C | Kabuli | Syria |
| FLIP_09_207C | Kabuli | Syria |
| FLIP_09_208C | Kabuli | Syria |
| FLIP_09_209C | Kabuli | Syria |
| FLIP_09_213C | Kabuli | Syria |
| FLIP_09_214C | Kabuli | Syria |
| FLIP_09_216C | Kabuli | Syria |
| FLIP_09_218C | Kabuli | Syria |
| FLIP_09_219C | Kabuli | Syria |
| FLIP_09_234C | Kabuli | Syria |
| FLIP_09_263C | Kabuli | Syria |
| FLIP_09_269C | Kabuli | Syria |
| FLIP_09_277C | Kabuli | Syria |
| FLIP_09_280C | Kabuli | Syria |
| FLIP_09_284C | Kabuli | Syria |
| FLIP_09_300C | Kabuli | Syria |
| FLIP_09_308C | Kabuli | Syria |
| FLIP_09_320C | Kabuli | Syria |
| FLIP_09_321C | Kabuli | Syria |
| FLIP_09_322C | Kabuli | Syria |
| FLIP_09_325C | Kabuli | Syria |
| FLIP_09_326C | Kabuli | Syria |
| FLIP_09_329C | Kabuli | Syria |
| FLIP_09_330C | Kabuli | Syria |
| FLIP_09_332C | Kabuli | Syria |
| FLIP_09_333C | Kabuli | Syria |
| FLIP_09_334C | Kabuli | Syria |
| FLIP_09_335C | Kabuli | Syria |
| FLIP_09_336C | Kabuli | Syria |
| FLIP_09_337C | Kabuli | Syria |
| FLIP_09_338C | Kabuli | Syria |
| FLIP_09_341C | Kabuli | Syria |
| FLIP_09_342C | Kabuli | Syria |
| FLIP_09_344C | Kabuli | Syria |
| FLIP_09_345C | Kabuli | Syria |
| FLIP_09_347C | Kabuli | Syria |
| FLIP_09_350C | Kabuli | Syria |
| FLIP_09_351C | Kabuli | Syria |
| FLIP_09_352C | Kabuli | Syria |
| FLIP_09_353C | Kabuli | Syria |
| FLIP_09_354C | Kabuli | Syria |
| FLIP_09_355C | Kabuli | Syria |
| FLIP_09_356C | Kabuli | Syria |
| FLIP_09_358C | Kabuli | Syria |
| FLIP_09_359C | Kabuli | Syria |
| FLIP_09_361C | Kabuli | Syria |
| FLIP_09_362C | Kabuli | Syria |
| FLIP_09_363C | Kabuli | Syria |
| FLIP_09_366C | Kabuli | Syria |
| FLIP_09_367C | Kabuli | Syria |
| FLIP_09_370C | Kabuli | Syria |
| FLIP_09_371C | Kabuli | Syria |
| FLIP_09_372C | Kabuli | Syria |
| FLIP_09_374C | Kabuli | Syria |
| FLIP_09_377C | Kabuli | Syria |
| FLIP_09_378C | Kabuli | Syria |
| FLIP_09_386C | Kabuli | Syria |
| FLIP_09_387C | Kabuli | Syria |
| FLIP_09_388C | Kabuli | Syria |
| FLIP_09_389C | Kabuli | Syria |
| FLIP_09_391C | Kabuli | Syria |
| FLIP_09_392C | Kabuli | Syria |
| FLIP_09_394C | Kabuli | Syria |
| FLIP_09_396C | Kabuli | Syria |
| FLIP_09_397C | Kabuli | Syria |
| FLIP_09_399C | Kabuli | Syria |
| FLIP_09_400C | Kabuli | Syria |
| FLIP_09_401C | Kabuli | Syria |
| FLIP_09_403C | Kabuli | Syria |
| FLIP_09_404C | Kabuli | Syria |
| FLIP_09_405C | Kabuli | Syria |
| FLIP_09_408C | Kabuli | Syria |
| FLIP_09_409C | Kabuli | Syria |
| FLIP_09_411C | Kabuli | Syria |
| FLIP_09_412C | Kabuli | Syria |
| FLIP_09_416C | Kabuli | Syria |
| FLIP_09_423C | Kabuli | Syria |
| FLIP_09_425C | Kabuli | Syria |
| FLIP_09_430C | Kabuli | Syria |
| FLIP_09_433C | Kabuli | Syria |
| FLIP_09_434C | Kabuli | Syria |
| FLIP_09_436C | Kabuli | Syria |
| FLIP_09_437C | Kabuli | Syria |
| FLIP_09_438C | Kabuli | Syria |
| FLIP_2001_56C | Kabuli | Syria |
| FLIP_2003_127C | Kabuli | Syria |
| FLIP_94_62C | Kabuli | Syria |
| FLIP_97_210C | Kabuli | Syria |
| FLIP_97_230C | Kabuli | Syria |
| GENESIS_079 | Kabuli | Australia |
| GENESIS_090 | Kabuli | Australia |
| GENESIS_KALKEE | Kabuli | Australia |
| ICC_01671 | Kabuli | India |
| ICC_06098 | Desi1 | India |
| ICC_11531 | Desi2 | India |
| ICC_14735 | Desi3 | India |
| ICC_16796 | Kabuli | India |
| ICCV_05314 | Desi4 | India |
| ICCV_91216 | Desi5 | India |
| ICCV_95224 | Desi6 | India |
| ICCV_96329 | Kabuli | India |
| IG_131610 | Kabuli | Kyrgyzstan |
| ILC_01272 | Desi7 | Turkey |
| KIMBERLEY_LARGE | Kabuli | Australia |
| KYABRA | Desi8 | Australia |
| NEC_02185C | Desi9 | India |
| PANT_G_114 | Desi10 | India |
| PBA_HATTRICK | Desi11 | Australia |
| PBA_MONARCH | Kabuli | Australia |
| PBA_PISTOL | Desi12 | Australia |
| PBA_SEAMER | Desi13 | Australia |
| PBA_SLASHER | Desi14 | Australia |
| S_98477 | Kabuli | India |

**Supplementary Figure 1.**
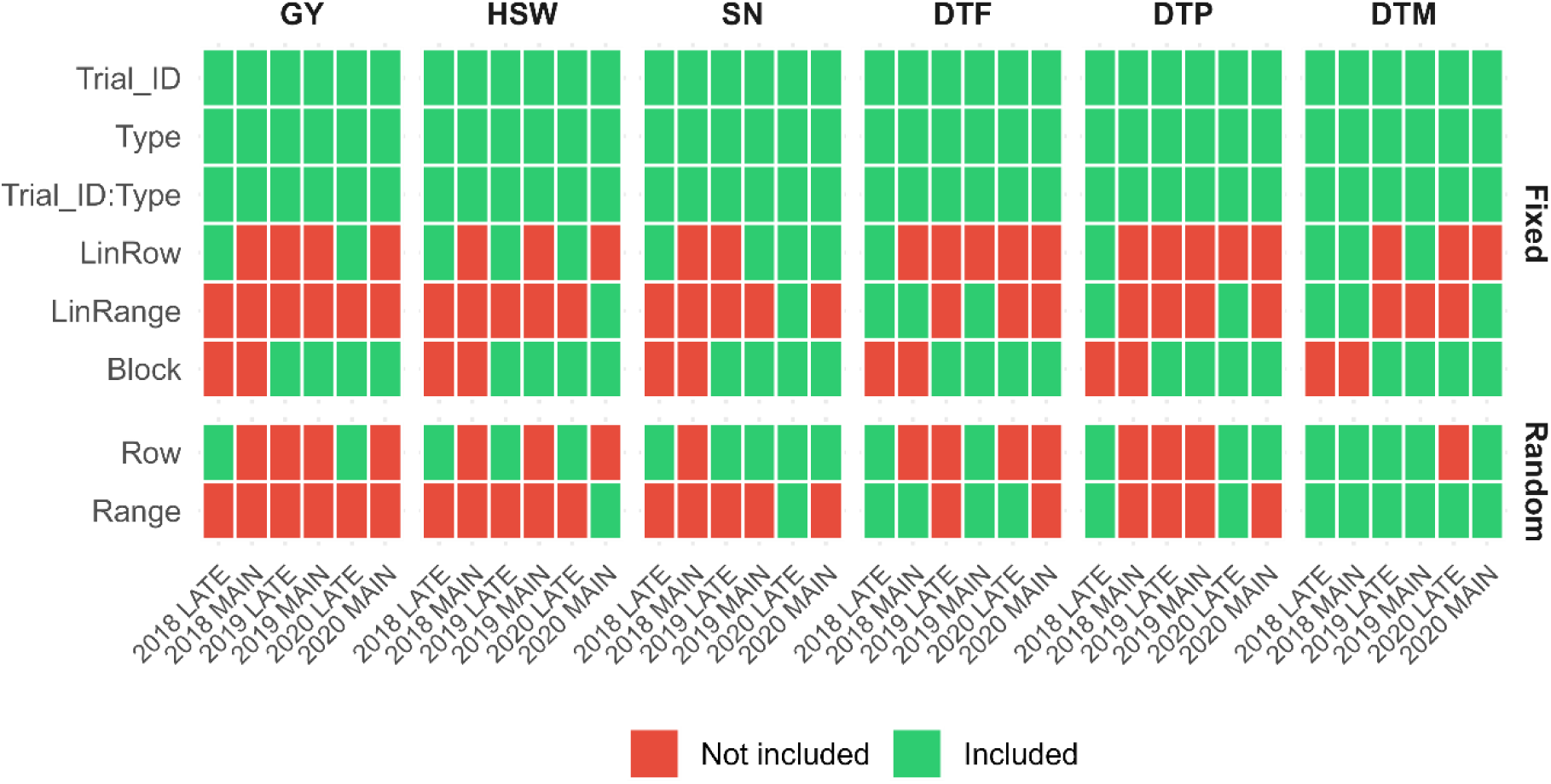
Fixed and random terms included in the multi environment trial (MET) model for each trait (GY, HSW, SN, DTF, DTP, DTM) across six trials at Narrabri from 2018 to 2020. Green indicates the term was included in the final model, red indicates it was not. All models included an autoregressive residual structure (AR1×AR1) within each trial.

**Supplementary Figure 2.**
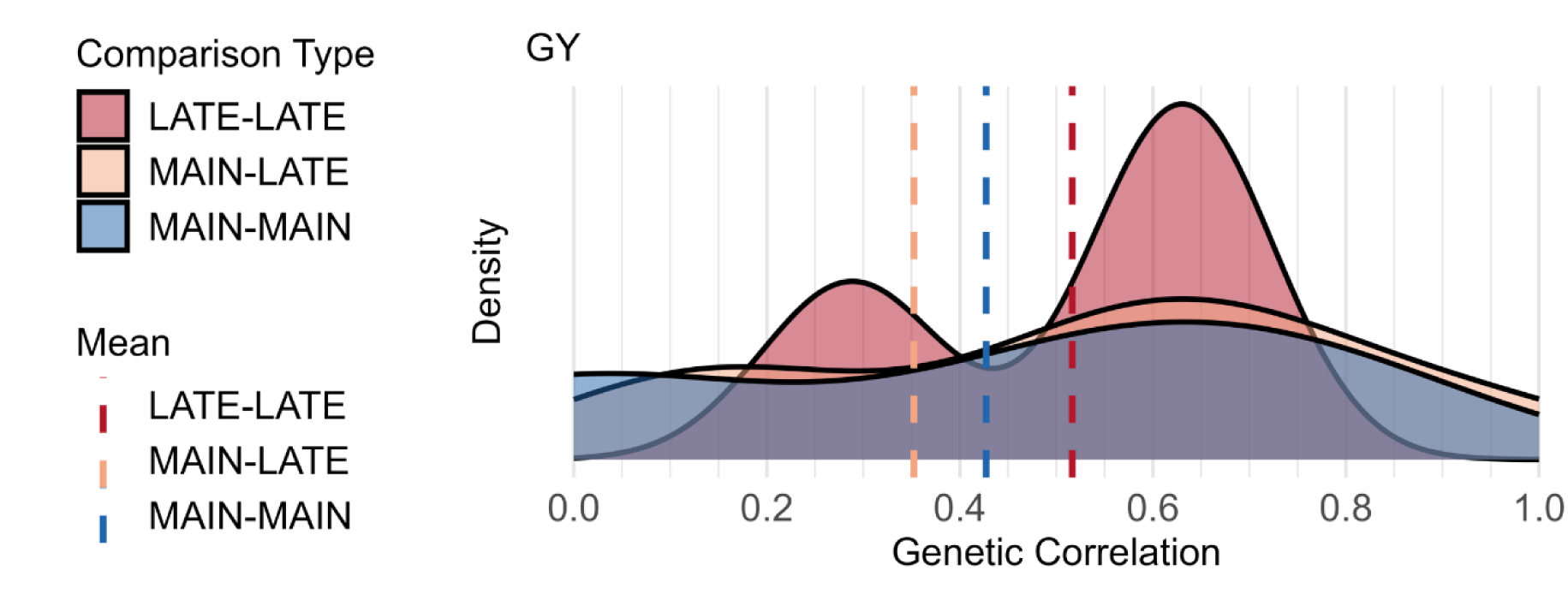
Density plot of pairwise genetic correlations from six trials at Narrabri from 2018 to 2020 for grain yield (GY). Grouped by time of sowing comparison: MAIN-MAIN (blue), LATE-LATE (red), and MAIN-LATE (orange). Dashed vertical lines indicate the mean genetic correlation for each group. Genetic correlations were derived from the additive genetic variance-covariance matrix estimated by the multi-environment trial (MET) model.

**Supplementary Table 2.** Factor loadings and variance partitioning parameters from the Factor Analytic 2 (FA2) model for grain yield (GY; t ha^-1^) that explained 84.5% of additive genetic variance.

| Trial | Factor 1<br>(additive) | Factor 2<br>(additive) | Factor 1<br>(nonadditive) | $\sigma^2G$ | $\sigma^2A\%$ | $H^2$ |
| --- | --- | --- | --- | --- | --- | --- |
| 2018 LATE | 0.08 | 0.00 | 0.03 | 0.03 | 0.34 | 0.70 |
| 2018 MAIN | 0.21 | -0.10 | 0.07 | 0.09 | 0.60 | 0.74 |
| 2019 LATE | 0.14 | 0.09 | 0.16 | 0.05 | 0.50 | 0.76 |
| 2019 MAIN | 0.11 | 0.03 | 0.19 | 0.06 | 0.23 | 0.60 |
| 2020 LATE | 0.06 | 0.09 | 0.10 | 0.03 | 0.70 | 0.64 |
| 2020 MAIN | 0.02 | -0.08 | 0.10 | 0.03 | 0.33 | 0.58 |

$\sigma^2G$ = total genetic variance; $\sigma^2A\%$ = proportion of $\sigma^2G$ that is additive; $H^2$ = broad-sense heritability estimated from DIAG model

**Supplementary Table 3.**
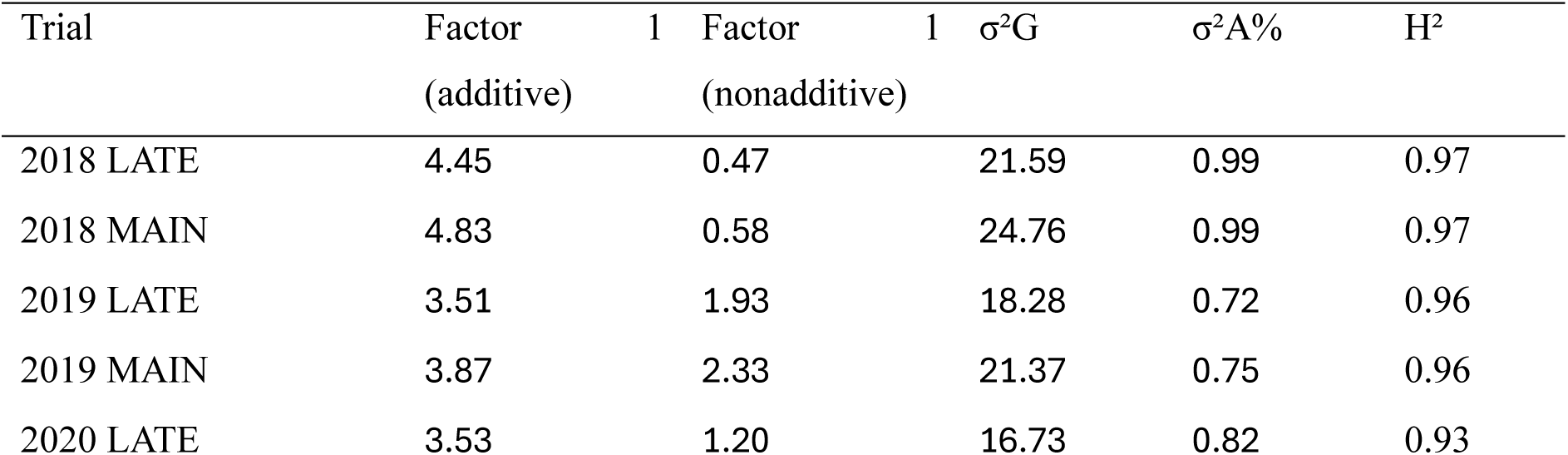
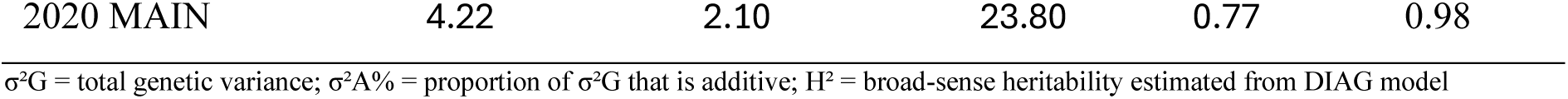
Factor loadings and variance partitioning parameters from the Factor Analytic 1 (FA1) model for 100 seed weight (HSW; g) that explained 94.2% of additive genetic variance.

**Supplementary Table 4.**
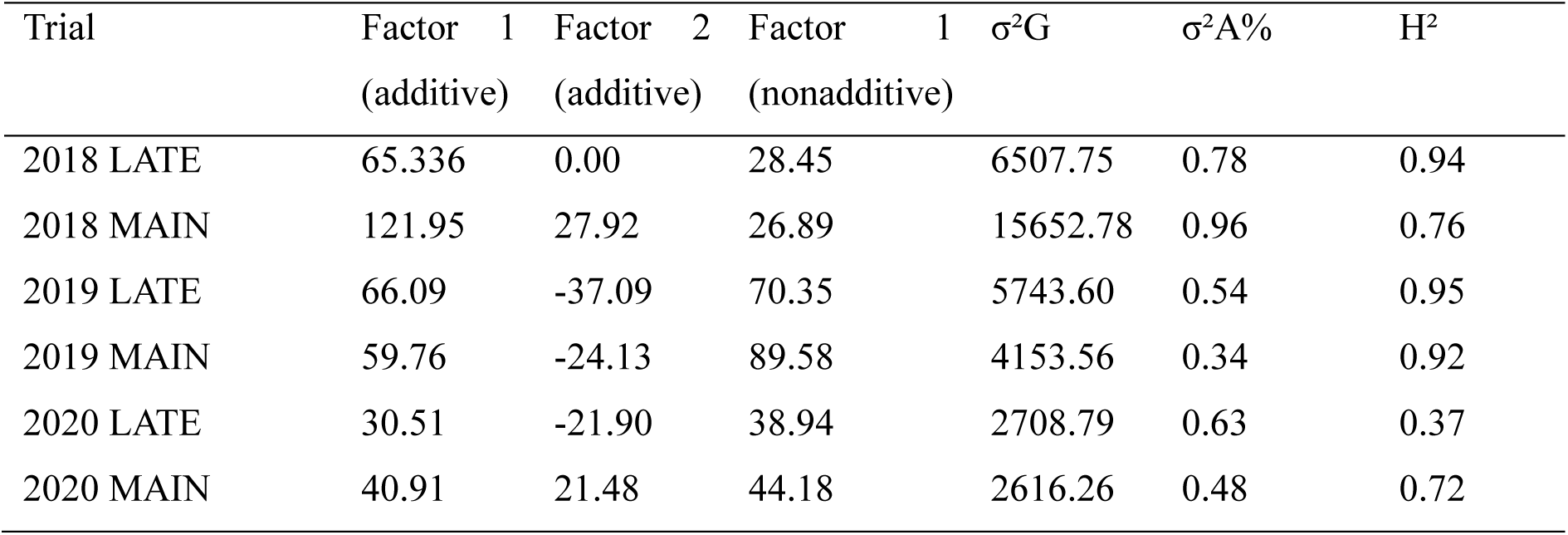
Factor loadings and variance partitioning parameters from the Factor Analytic 2 (FA2) model for seed number (SN; seeds m^2^) that explained 89.3% of additive genetic variance.

**Supplementary Table 5:** Factor loadings and variance partitioning parameters from the Factor Analytic 1 (FA1) model for thermal time to flowering (DTF; °Cd) that explained 98.3% of additive genetic variance.

| Trial | Factor 1<br>(additive) | Factor 1<br>(nonadditive) | $\sigma^2G$ | $\sigma^2A\%$ | $H^2$ |
| --- | --- | --- | --- | --- | --- |
| 2018 LATE | 122.19 | -1.33 | 16886.20 | 0.88 | 0.77 |
| 2018 MAIN | 50.78 | 5.60 | 2706.63 | 0.97 | 0.86 |
| 2019 LATE | 58.97 | 39.16 | 5180.84 | 0.70 | 0.95 |
| 2019 MAIN | 65.17 | 32.47 | 6080.79 | 0.70 | 0.91 |
| 2020 LATE | 64.52 | 8.59 | 4784.10 | 0.90 | 0.94 |
| 2020 MAIN | 42.27 | 2.01 | 2267.94 | 0.86 | 0.86 |

$\sigma^2G$ = total genetic variance; $\sigma^2A\%$ = proportion of $\sigma^2G$ that is additive; $H^2$ = broad-sense heritability estimated from DIAG model

**Supplementary Table 6:** Factor loadings and variance partitioning parameters from the Factor Analytic 1 (FA1) model for thermal time to podding (DTP; °Cd) that explained 96.6% of additive genetic variance.

| Trial | Factor 1<br>(additive) | Factor 1<br>(nonadditive) | $\sigma^2G$ | $\sigma^2A\%$ | $H^2$ |
| --- | --- | --- | --- | --- | --- |
| 2018 LATE | 34.07 | -7.89 | 1341.77 | 0.87 | 0.54 |
| 2018 MAIN | 51.90 | 8.94 | 3042.28 | 0.91 | 0.83 |
| 2019 LATE | 68.84 | 41.38 | 7133.40 | 0.68 | 0.94 |
| 2019 MAIN | 36.26 | 25.38 | 2119.94 | 0.62 | 0.77 |
| 2020 LATE | 55.05 | 13.35 | 3885.29 | 0.80 | 0.89 |
| 2020 MAIN | 39.93 | 11.49 | 2865.94 | 0.62 | 0.70 |

$\sigma^2G$ = total genetic variance; $\sigma^2A\%$ = proportion of $\sigma^2G$ that is additive; $H^2$ = broad-sense heritability estimated from DIAG model

**Supplementary Table 7:** Factor loadings and variance partitioning parameters from the Factor Analytic (FA3) model for thermal time to maturity (DTM; °Cd) that explained 86.5% of additive genetic variance.

| Trial | Factor 1<br>(additive) | Factor 2<br>(additive) | Factor 3<br>(additive) | Factor 1<br>(nonadditive) | $\sigma^2G$ | $\sigma^2A\%$ | $H^2$ |
| --- | --- | --- | --- | --- | --- | --- | --- |
| 2018 LATE | 39.69 | 0.00 | 0.00 | 13.74 | 1764.43 | 0.89 | 0.78 |
| 2018 MAIN | 24.67 | 14.23 | 0.00 | 0.85 | 933.20 | 0.87 | 0.19 |
| 2019 LATE | 22.76 | -0.17 | 23.28 | 25.79 | 1746.23 | 0.61 | 0.92 |
| 2019 MAIN | 13.31 | 6.53 | 18.87 | 24.34 | 1633.15 | 0.35 | 0.85 |
| 2020 LATE | 59.20 | -6.16 | 21.36 | 39.58 | 7712.62 | 0.64 | 0.75 |
| 2020 MAIN | 23.88 | 23.75 | -38.24 | 20.01 | 3739.25 | 0.89 | 0.60 |

$\sigma^2G$ = total genetic variance; $\sigma^2A\%$ = proportion of $\sigma^2G$ that is additive; $H^2$ = broad-sense heritability estimated from DIAG model

**Supplementary Table 8.** Haploblock parameters used for localGEBV calculation across eight chickpea chromosomes including, for each chromosome, the linkage disequilibrium (LD) threshold (r²), tolerance parameter (Tol), total number of markers, number of haploblocks identified, percentage of single-marker blocks, mean number of markers per block, and maximum number of markers per block are reported. Parameters were selected independently per chromosome to optimise block definition.

| Chromosome | LD<br>Threshold<br>($r^2$) | Tol | Number of<br>markers | Number<br>of blocks | % single<br>marker<br>blocks | Mean<br>markers<br>per block | Max<br>markers<br>per block |
| --- | --- | --- | --- | --- | --- | --- | --- |
| 1 | 0.50 | 1 | 893 | 286 | 60% | 3.12 | 36.0 |
| 2 | 0.40 | 1 | 1020 | 270 | 60% | 3.78 | 74.0 |
| 3 | 0.50 | 2 | 575 | 146 | 57% | 3.94 | 51.0 |
| 4 | 0.50 | 2 | 2870 | 640 | 61% | 4.48 | 123 |
| 5 | 0.40 | 2 | 588 | 156 | 62% | 3.77 | 41.0 |
| 6 | 0.60 | 2 | 897 | 285 | 61% | 3.15 | 37.0 |
| 7 | 0.40 | 2 | 958 | 247 | 61% | 3.88 | 60.0 |
| 8 | 0.60 | 1 | 292 | 79 | 53% | 3.70 | 59.0 |

**Supplementary Table 9.**
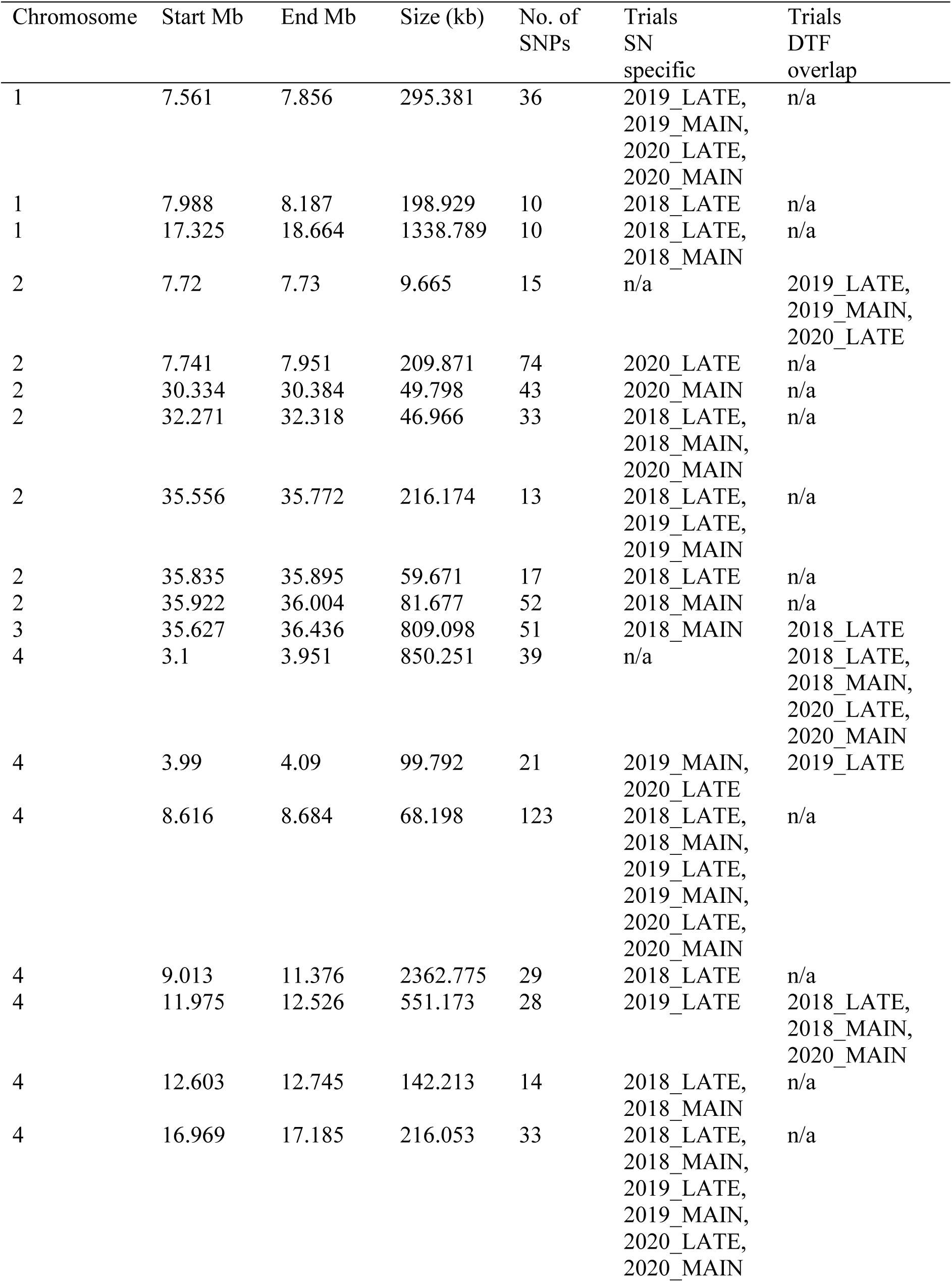
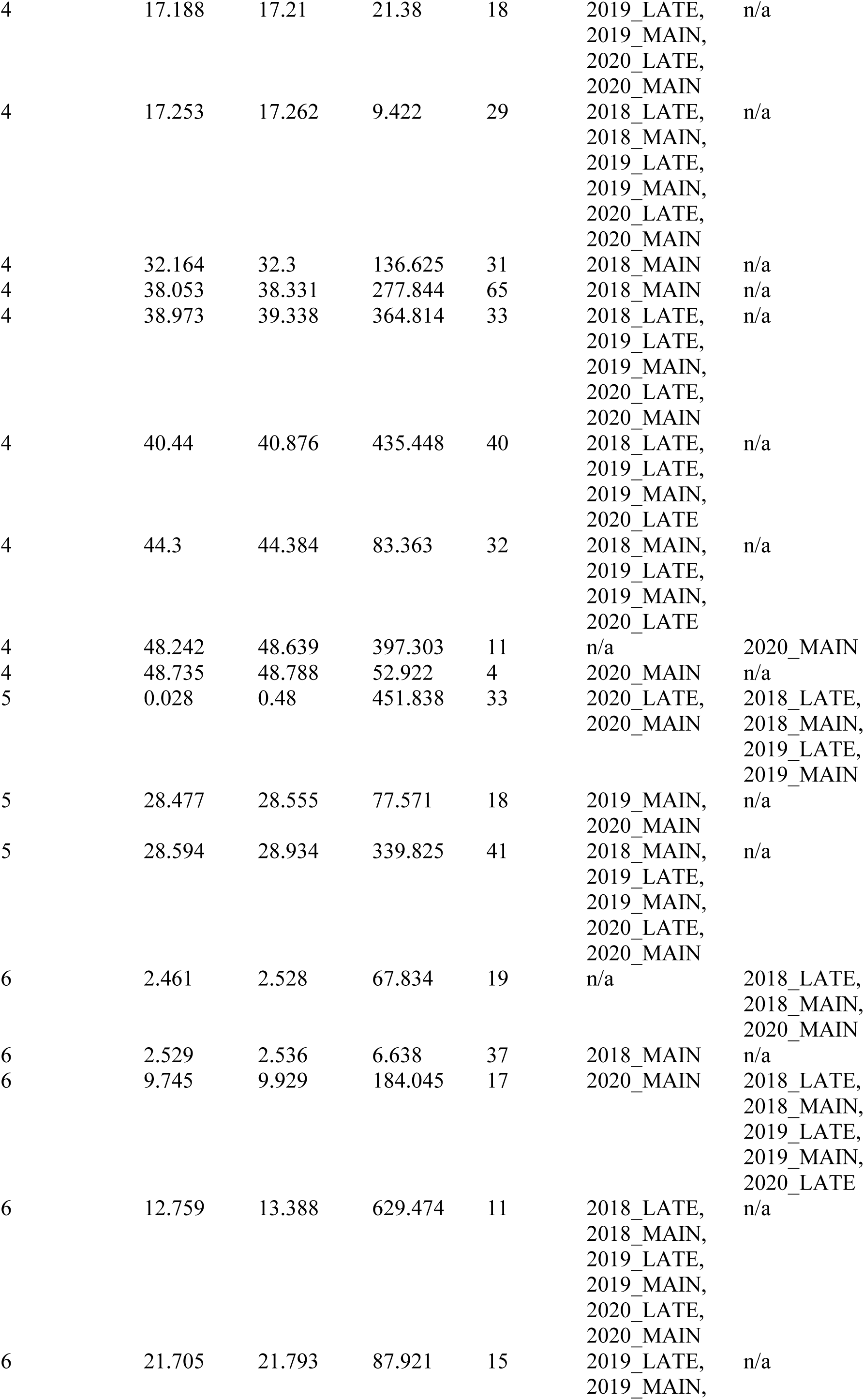
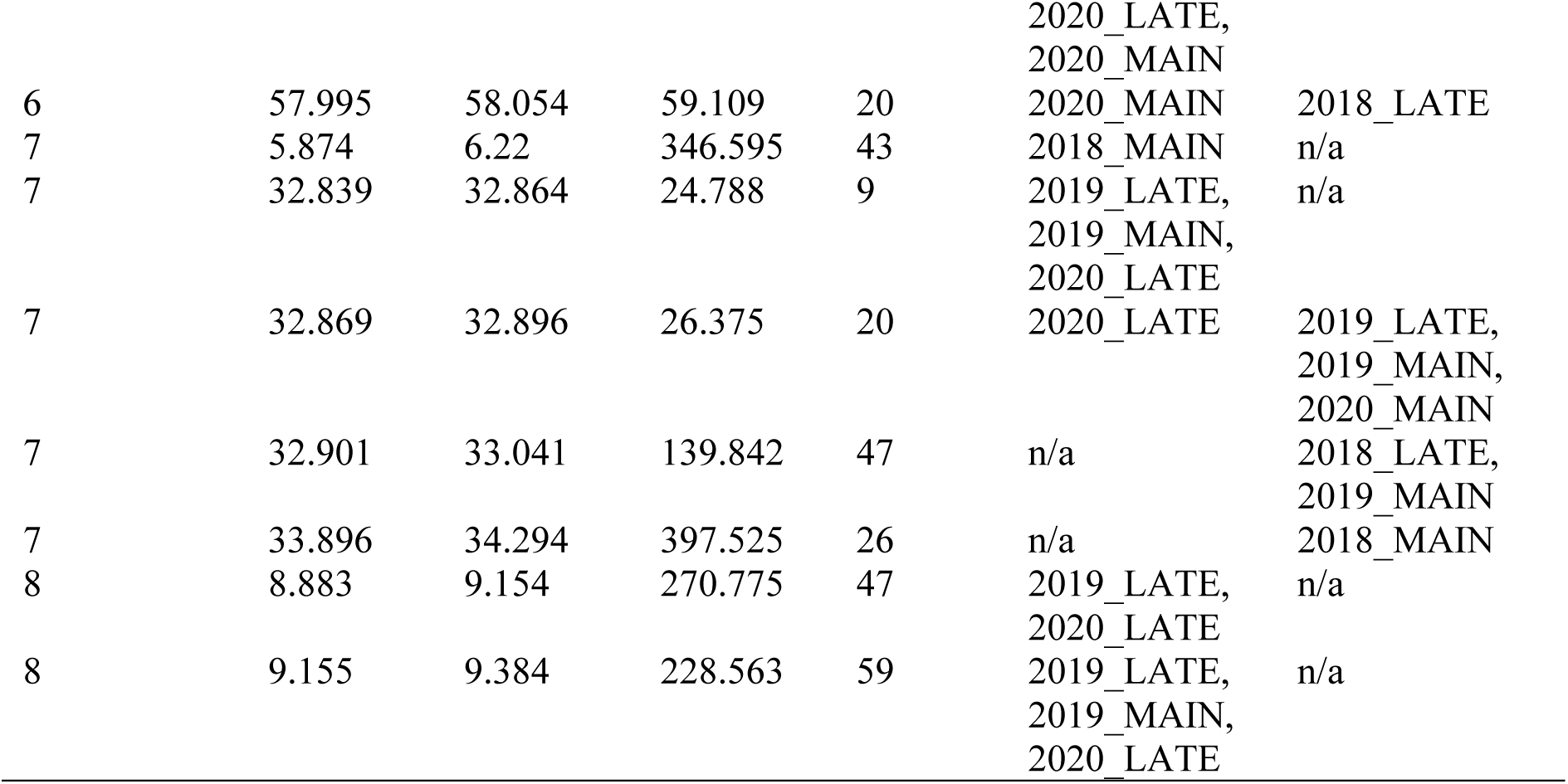
The top 1% of haploblocks with high block variance for seed number (SN) across six trials at Narrabri from 2018 to 2020, classified within each trial by overlap with the top 1% of haploblocks with high block variance for thermal time to flowering (DTF). For each block, the table reports the chromosome (Chr), genomic coordinates, size, number of SNPs, and the trials in which the block was classified as SN-specific or DTF-overlapping.

**Supplementary Figure 3.**
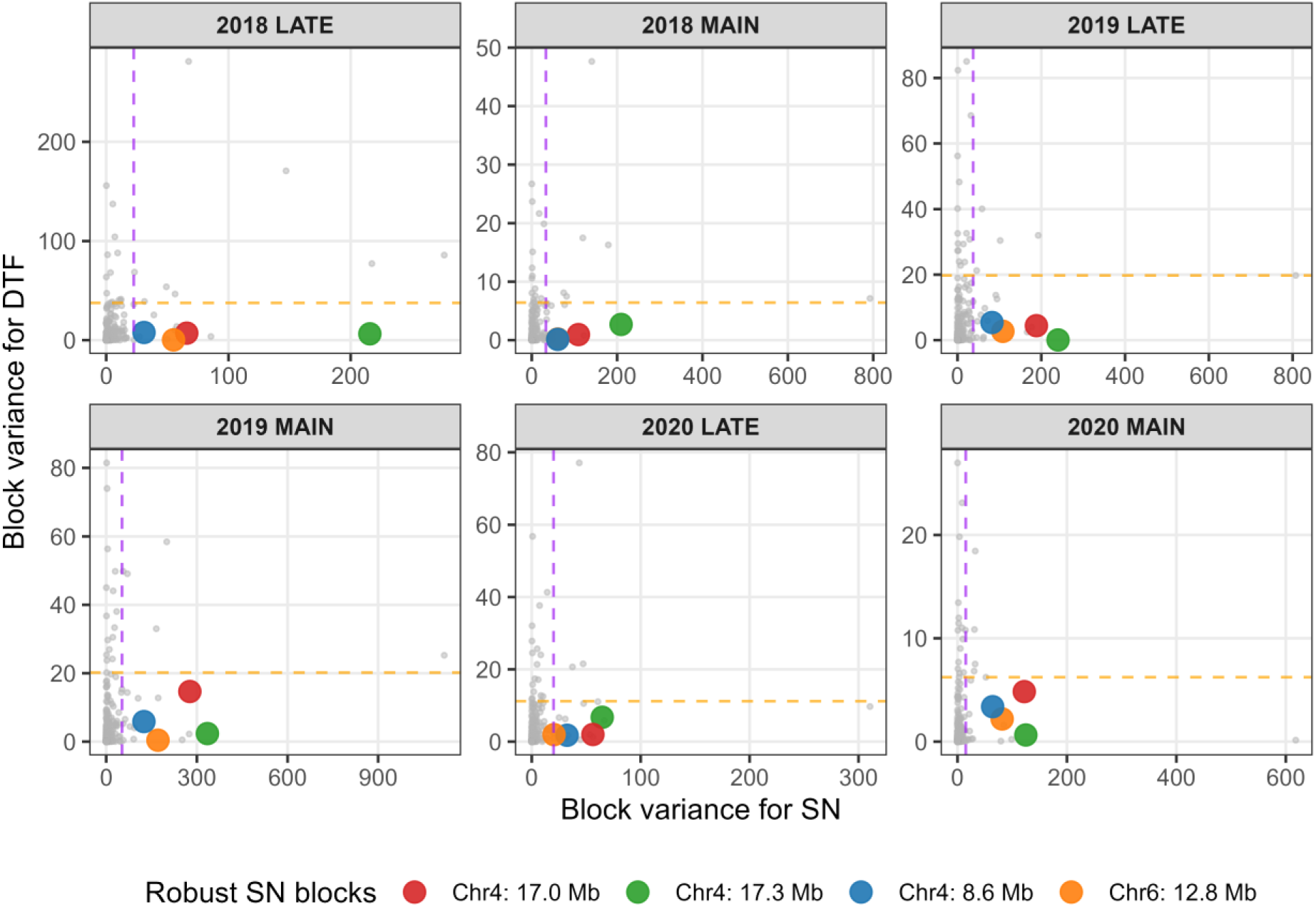
Block variance for seed number (SN) and thermal time to flowering (DTF) for all 2,109 haploblocks across six trials at Narrabri from 2018 to 2020. Each point represents a single haploblock, coloured points highlight the four haploblocks identified in the top 1% for SN and not the top 1% for DTF in all six trials. Dashed lines indicate the 1% threshold (rank 21 of 2,109) for SN (purple) and DTF (orange) within each trial.

**Supplementary Figure 4.**
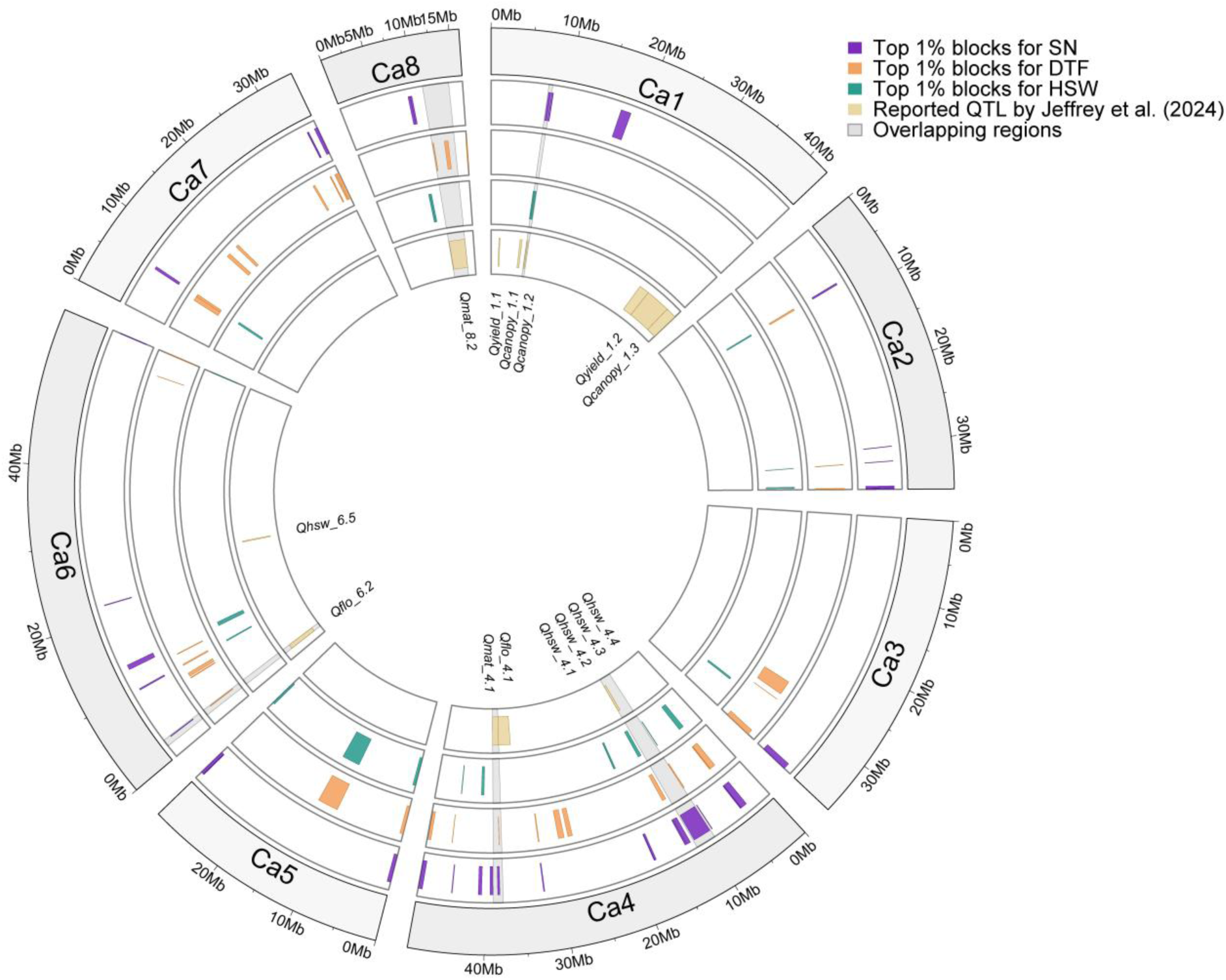
Genome-wide Circos plot comparing the physical positions of the top 1% blocks identified using the local GEBV approach in the present study for Seed Number (SN), Days to Flowering (DTF), and 100-seed weight (HSW) with the QTL reported by Jeffrey et al. (2024). From the outer to inner circles, tracks represent identified blocks for SN, DTF, HSW, and previously reported QTL from Jeffrey et al. (2024), respectively. Shaded regions indicate overlapping blocks identified in the present study and reported QTL.

## References

Aldiss, Z., Lam, Y., Baraibar, S., Van Der Meer, S., Dinglasan, E., Massel, K., Crisp, P., Godwin, I., Borrell, A., Moody, D., Hickey, L., & Robinson, H. (2025). Haplotype-based insights into seminal root angle in barley. The Plant Genome, 18(3), e70088. 10.1002/tpg2.70088

Anwar, M. R., Luckett, D. J., Chauhan, Y. S., Ip, R. H. L., Maphosa, L., Simpson, M., Warren, A., Raman, R., Richards, M. F., Pengilley, G., Hobson, K., & Graham, N. (2022). Modelling the effects of cold temperature during the reproductive stage on the yield of chickpea (Cicer arietinum L.). Int J Biometeorol, 66(1), 111–125. 10.1007/s00484-021-02197-8

Arriagada, O., Cacciuttolo, F., Cabeza, R. A., Carrasco, B., & Schwember, A. R. (2022). A Comprehensive Review on Chickpea (Cicer arietinum L.) Breeding for Abiotic Stress Tolerance and Climate Change Resilience. Int J Mol Sci, 23(12). 10.3390/ijms23126794

Baker, R. (1988). Tests for crossover genotype-environmental interactions. Canadian journal of plant science, 68(2), 405–410.

Barmukh, R., Roorkiwal, M., Garg, V., Khan, A. W., German, L., Jaganathan, D., Chitikineni, A., Kholova, J., Kudapa, H., Sivasakthi, K., Samineni, S., Kale, S. M., Gaur, P. M., Sagurthi, S. R., Benitez-Alfonso, Y., & Varshney, R. K. (2022). Genetic variation in CaTIFY4b contributes to drought adaptation in chickpea. Plant Biotechnol J, 20(9), 1701–1715. 10.1111/pbi.13840

Berger, J., Turner, N., Siddique, K., Knights, E., Brinsmead, R., Mock, I., Edmondson, C., & Khan, T. (2004). Genotype by environment studies across Australia reveal the importance of phenology for chickpea (Cicer arietinum L.) improvement. Australian Journal of Agricultural Research, 55(10), 1071–1084.

Bonhomme, R. (2000). Bases and limits to using ‘degree.day’ units. European Journal of Agronomy, 13(1), 1–10. 10.1016/S1161-0301(00)00058-7

Butler, D. G., Cullis, B. R., Gilmour, A. R., Gogel, B. G., & Thompson, R. (2023). ASReml-R Reference Manual Version 4.2. In VSN International Ltd.

Chang, C. C., Chow, C. C., Tellier, L. C., Vattikuti, S., Purcell, S. M., & Lee, J. J. (2015). Second-generation PLINK: rising to the challenge of larger and richer datasets. Gigascience, 4, 7. 10.1186/s13742-015-0047-8

Chauhan, Y., Allard, S., Williams, R., Williams, B., Mundree, S., Chenu, K., & Rachaputi, N. C. (2017). Characterisation of chickpea cropping systems in Australia for major abiotic production constraints. Field Crops Research, 204, 120–134. 10.1016/j.fcr.2017.01.008

Chauhan, Y. S., Ryan, M., Chandra, S., & Sadras, V. O. (2019). Accounting for soil moisture improves prediction of flowering time in chickpea and wheat. Sci Rep, 9(1), 7510. 10.1038/s41598-019-43848-6

Costa-Neto, G., Galli, G., Carvalho, H. F., Crossa, J., & Fritsche-Neto, R. (2021). EnvRtype: a software to interplay enviromics and quantitative genomics in agriculture. G3 (Bethesda), 11(4). 10.1093/g3journal/jkab040

Cullis, B. R., Smith, A. B., & Coombes, N. E. (2006). On the design of early generation variety trials with correlated data. Journal of agricultural, biological, and environmental statistics, 11(4), 381–393.

Danakumara, T., Kumar, N., Patil, B. S., Kumar, T., Bharadwaj, C., Jain, P. K., Nimmy, M. S., Joshi, N., Parida, S. K., Bindra, S., Kole, C., & Varshney, R. K. (2024). Unraveling the genetics of heat tolerance in chickpea landraces (Cicer arietinum L.) using genome-wide association studies. Frontiers in Plant Science, 15. 10.3389/fpls.2024.1376381

Das, S., Forer, L., Schönherr, S., Sidore, C., Locke, A. E., Kwong, A., Vrieze, S. I., Chew, E. Y., Levy, S., McGue, M., Schlessinger, D., Stambolian, D., Loh, P. R., Iacono, W. G., Swaroop, A., Scott, L. J., Cucca, F., Kronenberg, F., Boehnke, M.,…Fuchsberger, C. (2016). Next-generation genotype imputation service and methods. Nat Genet, 48(10), 1284–1287. 10.1038/ng.3656

Davies, S., Turner, N., Siddique, K., Leport, L., & Plummer, J. (1999). Seed growth of desi and kabuli chickpea (Cicer arietinum L.) in a short-season Mediterranean-type environment. Australian Journal of Experimental Agriculture, 39(2), 181–188.

Devasirvatham, V., Gaur, P. M., Raju, T. N., Trethowan, R. M., & Tan, D. K. Y. (2015a). Field response of chickpea (Cicer arietinum L.) to high temperature. Field Crops Research, 172, 59–71. 10.1016/j.fcr.2014.11.017

Devasirvatham, V., Tan, D., Gaur, P., & Trethowan, R. (2015b). Chickpea and temperature stress: An overview. In (pp. 81-90). 10.1002/9781118917091.ch5

Devasirvatham, V., Tan, D. K. Y., Gaur, P. M., Raju, T. N., & Trethowan, R. M. (2012). High temperature tolerance in chickpea and its implications for plant improvement. CROP & PASTURE SCIENCE, 63(5), 419–428. 10.1071/cp11218

Dreccer, M. F., Fainges, J., Whish, J., Ogbonnaya, F. C., & Sadras, V. O. (2018). Comparison of sensitive stages of wheat, barley, canola, chickpea and field pea to temperature and water stress across Australia. Agricultural and Forest Meteorology, 248, 275–294. 10.1016/j.agrformet.2017.10.006

Fang, X., Turner, N. C., Yan, G., Li, F., & Siddique, K. H. M. (2010). Flower numbers, pod production, pollen viability, and pistil function are reduced and flower and pod abortion increased in chickpea (Cicer arietinum L.) under terminal drought. Journal of Experimental Botany, 61(2), 335–345. 10.1093/jxb/erp307

FAO. (2023). https://www.fao.org/faostat/en/#data/QCL

Fischer, R. (2016). The effect of duration of the vegetative phase in irrigated semi-dwarf spring wheat on phenology, growth and potential yield across sowing dates at low latitude. Field Crops Research, 198, 188–199.

Garg, V., Barmukh, R., Huang, Y., Chitikineni, A., Hobson, K., Yang, B., Jia, Y., Bi, S., Kaur, S., Asif, M. A., Hayden, M., Norton, S., Sharma, D. L., Siddique, K. H. M., Liu, X., Li, C., & Varshney, R. K. (2025). An Australian chickpea pan-genome provides insights into genome organization and offers opportunities for enhancing drought adaptation for crop improvement. Plant Biotechnology Journal, n/a(n/a). 10.1111/pbi.70192

Gilmour, A. R., Cullis, B. R., & Verbyla, A. (1997). Accounting for Natural and Extraneous Variation in the Analysis of Field Experiments. Journal of agricultural, biological, and environmental statistics, 2, 269–293.

Gimenez, R., Lake, L., Cossani, C. M., Ortega Martinez, R., Hayes, J. E., Dreccer, M. F., French, R., Weller, J. L., & Sadras, V. O. (2025). Linking phenology, harvest index, and genetics to improve chickpea grain yield. J Exp Bot, 76(6), 1658–1677. 10.1093/jxb/erae487

GRDC. (2016). Chickpeas Southern Region - GrowNotes™. https://grdc.com.au/resources-and-publications/grownotes/crop-agronomy/chickpea-southern-region-grownotes

Hijmans, R. (2024). geosphere: Spherical Trigonometry. In https://cran.r-project.org/web/packages/geosphere/index.html

Hiremath, P. J., Farmer, A., Cannon, S. B., Woodward, J., Kudapa, H., Tuteja, R., Kumar, A., BhanuPrakash, A., Mulaosmanovic, B., Gujaria, N., Krishnamurthy, L., Gaur, P. M., KaviKishor, P. B., Shah, T., Srinivasan, R., Lohse, M., Xiao, Y., Town, C. D., Cook, D. R.,…Varshney, R. K. (2011). Large-scale transcriptome analysis in chickpea (Cicer arietinum L.), an orphan legume crop of the semi-arid tropics of Asia and Africa. Plant Biotechnology Journal, 9(8), 922–931. 10.1111/j.1467-7652.2011.00625.x

Hobson, K., Dron, N., Day, S., Borgognone, M. G., & McMurray, L. (2016). Developing medium to large seeded kabuli chickpeas with early maturity, improved yield and Ascochyta bight resistance for Australian growers. Australian Pulse Conference, Tamworth, NSW.

Incitec Pivot Fertiliers. https://www.incitecpivotfertilisers.com.au/products-services/our-products/compounds/granulock-z/

Jarquín, D., Crossa, J., Lacaze, X., Du Cheyron, P., Daucourt, J., Lorgeou, J., Piraux, F., Guerreiro, L., Pérez, P., Calus, M., Burgueño, J., & de los Campos, G. (2014). A reaction norm model for genomic selection using high-dimensional genomic and environmental data. Theor Appl Genet, 127(3), 595–607. 10.1007/s00122-013-2243-1

Jeffrey, C., Kaiser, B., Trethowan, R., & Ziems, L. (2024). Genome-wide association study reveals heat tolerance QTL for canopy-closure and early flowering in chickpea [Original Research]. Frontiers in Plant Science, Volume 15 - 2024. 10.3389/fpls.2024.1458250

Jeffrey, C., Ziems, L., Kaiser, B., & Trethowan, R. (2025). A growing degree day model determines the effect of temperature stress on diverse chickpea genotypes. Frontiers in Plant Science, 15. 10.3389/fpls.2024.1496629

Jha, U. C., Nath, C. P., Paul, P. J., Nayyar, H., Kumar, N., Dixit, G. P., Sen, S., Kumar, Y., & Prasad, P. V. V. (2025). Decoding the heat stress resilience in Chickpea (Cicer arietinum L.): multi-trait analysis for genotypic adaptation. Scientific Reports, 15(1), 25055. 10.1038/s41598-025-07573-7

Jha, U. C., Nayyar, H., Thudi, M., Beena, R., Vara Prasad, P. V., & Siddique, K. H. M. (2024). Unlocking the nutritional potential of chickpea: strategies for biofortification and enhanced multinutrient quality [Review]. Frontiers in Plant Science, Volume 15 - 2024. 10.3389/fpls.2024.1391496

Kale, S. M., Jaganathan, D., Ruperao, P., Chen, C., Punna, R., Kudapa, H., Thudi, M., Roorkiwal, M., Katta, M. A., & Doddamani, D. (2015). Prioritization of candidate genes in “QTL-hotspot” region for drought tolerance in chickpea (Cicer arietinum L.). Scientific Reports, 5(1), 15296.

Kaushal, N., Awasthi, R., Gupta, K., Gaur, P., Siddique, K. H. M., & Nayyar, H. (2013). Heat-stress-induced reproductive failures in chickpea (Cicer arietinum) are associated with impaired sucrose metabolism in leaves and anthers. Funct Plant Biol, 40(12), 1334–1349. 10.1071/fp13082

Kirkpatrick, M., & Meyer, K. (2004). Direct estimation of genetic principal components: simplified analysis of complex phenotypes. Genetics, 168(4), 2295–2306.

Krishnamurthy, L., Gaur, P., Basu, P., Chaturvedi, S., Tripathi, S., Vadez, V., Rathore, A., Varshney, R., & Gowda, C. (2011). Large genetic variation for heat tolerance in the reference collection of chickpea (Cicer arietinum L.) germplasm. Plant Genetic Resources, 9(1), 59–69.

Lake, L., Manson, J. B., Severini, A. D., Chauhan, Y., Chenu, K., Smith, M. R., & Sadras, V. O. (2025). Spatial probabilistic patterns of joint heat and water stress of chickpea in Australia. bioRxiv, 2025.2007.2009.664009. 10.1101/2025.07.09.664009

Lake, L., & Sadras, V. O. (2014). The critical period for yield determination in chickpea (Cicer arietinum L.). Field Crops Research, 168, 1–7. 10.1016/j.fcr.2014.08.003

Lake, L., & Sadras, V. O. (2016). Screening chickpea for adaptation to water stress: Associations between yield and crop growth rate. European Journal of Agronomy, 81, 86–91.

Leport, L., Turner, N. C., Davies, S. L., & Siddique, K. H. M. (2006). Variation in pod production and abortion among chickpea cultivars under terminal drought. European Journal of Agronomy, 24(3), 236–246. 10.1016/j.eja.2005.08.005

Li, Y., Lake, L., Chauhan, Y. S., Taylor, J., & Sadras, V. O. (2022). Genetic basis and adaptive implications of temperature-dependent and temperature-independent effects of drought on chickpea reproductive phenology. J Exp Bot, 73(14), 4981–4995. 10.1093/jxb/erac195

Masson-Delmotte, V., Zhai, P., Pirani, A., Connors, S. L., Péan, C., Berger, S., Caud, N., Chen, Y., Goldfarb, L., & Gomis, M. (2021). Climate change 2021: the physical science basis. Contribution of working group I to the sixth assessment report of the intergovernmental panel on climate change, 2(1), 2391.

McMaster, G. S., & Wilhelm, W. W. (1997). Growing degree-days: one equation, two interpretations. Agricultural and Forest Meteorology, 87(4), 291–300. 10.1016/S0168-1923(97)00027-0

Meek, D. W., Hatfield, J. L., Howell, T. A., Idso, S. B., & Reginato, R. J. (1984). A Generalized Relationship between Photosynthetically Active Radiation and Solar Radiation. Agronomy Journal, 76(6), 939–945. 10.2134/agronj1984.00021962007600060018x

Merga, B., & Haji, J. (2019). Economic importance of chickpea: Production, value, and world trade. Cogent Food & Agriculture, 5(1), 1615718. 10.1080/23311932.2019.1615718

Nakano, H., Kobayashi, M., & Terauchi, T. (1998). Sensitive Stages to Heat Stress in Pod Setting of Common Bean (Phaseolus vulgaris L.). Japanese Journal of Tropical Agriculture, 42(2), 78–84. 10.11248/jsta1957.42.78

Nguyen, D. T., Hayes, J. E., Harris, J., & Sutton, T. (2022). Fine Mapping of a Vigor QTL in Chickpea (Cicer arietinum L.) Reveals a Potential Role for Ca4_TIFY4B in Regulating Leaf and Seed Size [Original Research]. Frontiers in Plant Science, *Volume 13 -* 2022. 10.3389/fpls.2022.829566

Nix, H. (1976). Climate and crop productivity. Climate and Rice. International Rice Research Institute (IRRI), Los Bafios, 495.

Novozymes. https://biosolutions.novozymes.com/sites/default/files/field_media_document/2021-11/TagTeam%20CA%20EN.pdf.

Nufarm. https://nufarm.com/au/product/p-pickel-t/

Piepho, H.-P., & Blancon, J. (2023). Extending Finlay–Wilkinson regression with environmental covariates. Plant Breeding, 142(5), 621–631. 10.1111/pbr.13130

Piepho, H. P., Möhring, J., Melchinger, A. E., & Büchse, A. (2008). BLUP for phenotypic selection in plant breeding and variety testing. Euphytica, 161(1), 209–228. 10.1007/s10681-007-9449-8

Purcell, S., & Chang, C. (2025). PLINK [1.9]. www.cog-genomics.org/plink/1.9/

Pushpavalli, R., Zaman-Allah, M., Turner, N. C., Baddam, R., Rao, M. V., & Vadez, V. (2015). Higher flower and seed number leads to higher yield under water stress conditions imposed during reproduction in chickpea. Funct Plant Biol, 42(2), 162–174. 10.1071/fp14135

R Core Team, R. (2025). R: A language and environment for statistical computing. In: Citeseer.

Rani, A., Devi, P., Jha, U. C., Sharma, K. D., Siddique, K. H., & Nayyar, H. (2020). Developing climate-resilient chickpea involving physiological and molecular approaches with a focus on temperature and drought stresses. Frontiers in Plant Science, 10, 1759.

Robertson, M., Carberry, P., Huth, N., Turpin, J., Probert, M., Poulton, P., Bell, M., Wright, G., Yeates, S., & Brinsmead, R. (2002). Simulation of growth and development of diverse legume species in APSIM. Australian Journal of Agricultural Research, 53, 429–446. 10.1071/AR01106

Rodriguez, D., & Sadras, V. (2007). The limit to wheat water-use efficiency in eastern Australia. I.* Gradients in the radiation environment and atmospheric demand. Australian Journal of Agricultural Research, 58(4), 287–302.

Rogers, J. S. (1972). Measures of genetic similarity and genetic distance. Studies in genetics VII, 145-153.

Sadras, V. O., Vadez, V., Purushothaman, R., Lake, L., & Marrou, H. (2015). Unscrambling confounded effects of sowing date trials to screen for crop adaptation to high temperature. Field Crops Research, 177, 1–8. 10.1016/j.fcr.2015.02.024

Sandana, P., & Calderini, D. F. (2012). Comparative assessment of the critical period for grain yield determination of narrow-leafed lupin and pea. European Journal of Agronomy, 40, 94–101.

Shaffer, W., Papin, V., Yadav, S., Voss-Fels, K. P., Hickey, L. T., Hayes, B. J., & Dinglasan, E. G. (2025). Local genomic estimates provide a powerful framework for haplotype discovery. bioRxiv, 2025.2008.2028.672830. 10.1101/2025.08.28.672830

Siddique, K., & Regan, K. (2005). Registration of ‘Kimberley Large’ Kabuli Chickpea. Crop Science, 45, 1659–1660. 10.2135/cropsci2004.0405

Siddique, K. H. M., Regan, K. L., & Malhotra, R. S. (2007). Registration of ‘Almaz’ Kabuli Chickpea Cultivar. Crop Science, 47(1), 437–437. 10.2135/cropsci2006.02.0114

Smith, A., Cullis, B., & Thompson, R. (2004). Analyzing Variety by Environment Data Using Multiplicative Mixed Models and Adjustments for Spatial Field Trend. Biometrics, 57(4), 1138–1147. 10.1111/j.0006-341X.2001.01138.x

Smith, A. B., & Cullis, B. R. (2018). Plant breeding selection tools built on factor analytic mixed models for multi-environment trial data. Euphytica, 214(8), 143. 10.1007/s10681-018-2220-5

Soltani, A., Hammer, G. L., Torabi, B., Robertson, M. J., & Zeinali, E. (2006). Modeling chickpea growth and development: Phenological development. Field Crops Research, 99(1), 1–13. 10.1016/j.fcr.2006.02.004

Soltani, A., & Sinclair, T. R. (2011). A simple model for chickpea development, growth and yield. Field Crops Research, 124(2), 252–260. 10.1016/j.fcr.2011.06.021

Stella, A. A., Pavan, J. P. S., Araújo, M. S., Fregonezi, B. F., Unzimai, I. V., Leles, E. P., Santos, M. F., Goldsmith, P., Chigeza, G., Diers, B. W., Gathungu, T., Njoroge, J., & Pinheiro, J. B. (2025). Soybean selection in Kenya enhanced by multi-trait and genotype-by-environment interaction modeling. Scientific Reports, 15(1), 27575. 10.1038/s41598-025-10654-2

Subedi, M., Naiker, M., du Preez, R., Adorada, D. L., & Bhattarai, S. (2024). Evaluation of Kabuli Chickpea Genotypes for Tropical Adaptation in Northern Australia. Agriculture, 14(10), 1851. https://www.mdpi.com/2077-0472/14/10/1851

Tardieu, F., & Tuberosa, R. (2010). Dissection and modelling of abiotic stress tolerance in plants. Current Opinion in Plant Biology, 13(2), 206–212. 10.1016/j.pbi.2009.12.012

VanRaden, P. M. (2008). Efficient Methods to Compute Genomic Predictions. Journal of Dairy Science, 91(11), 4414–4423. 10.3168/jds.2007-0980

Varshney, R. K., Hiremath, P. J., Lekha, P., Kashiwagi, J., Balaji, J., Deokar, A. A., Vadez, V., Xiao, Y., Srinivasan, R., Gaur, P. M., Siddique, K. H. M., Town, C. D., & Hoisington, D. A. (2009). A comprehensive resource of drought- and salinity- responsive ESTs for gene discovery and marker development in chickpea (Cicer arietinum L.). BMC genomics, 10(1), 523. 10.1186/1471-2164-10-523

Varshney, R. K., Song, C., Saxena, R. K., Azam, S., Yu, S., Sharpe, A. G., Cannon, S., Baek, J., Rosen, B. D., Tar’an, B., Millan, T., Zhang, X., Ramsay, L. D., Iwata, A., Wang, Y., Nelson, W., Farmer, A. D., Gaur, P. M., Soderlund, C.,…Cook, D. R. (2013). Draft genome sequence of chickpea (Cicer arietinum) provides a resource for trait improvement. Nature Biotechnology, 31(3), 240–246. 10.1038/nbt.2491

Varshney, R. K., Thudi, M., Nayak, S. N., Gaur, P. M., Kashiwagi, J., Krishnamurthy, L., Jaganathan, D., Koppolu, J., Bohra, A., Tripathi, S., Rathore, A., Jukanti, A. K., Jayalakshmi, V., Vemula, A., Singh, S. J., Yasin, M., Sheshshayee, M. S., & Viswanatha, K. P. (2014). Genetic dissection of drought tolerance in chickpea (Cicer arietinum L.). Theoretical and Applied Genetics, 127(2), 445–462. 10.1007/s00122-013-2230-6

Varshney, R. K., Thudi, M., Roorkiwal, M., He, W., Upadhyaya, H. D., Yang, W., Bajaj, P., Cubry, P., Rathore, A., Jian, J., Doddamani, D., Khan, A. W., Garg, V., Chitikineni, A., Xu, D., Gaur, P. M., Singh, N. P., Chaturvedi, S. K., Nadigatla, G. V. P. R.,…Liu, X. (2019). Resequencing of 429 chickpea accessions from 45 countries provides insights into genome diversity, domestication and agronomic traits. Nature Genetics, 51(5), 857–864. 10.1038/s41588-019-0401-3

Voss-Fels, K. P., Stahl, A., Wittkop, B., Lichthardt, C., Nagler, S., Rose, T., Chen, T.-W., Zetzsche, H., Seddig, S., Majid Baig, M., Ballvora, A., Frisch, M., Ross, E., Hayes, B. J., Hayden, M. J., Ordon, F., Leon, J., Kage, H., Friedt, W.,…Snowdon, R. J. (2019). Breeding improves wheat productivity under contrasting agrochemical input levels. Nature Plants, 5(7), 706–714. 10.1038/s41477-019-0445-5

Weffort, V. R. S., & Lamounier, J. A. (2024). Hidden hunger - a narrative review. J Pediatr (Rio J), 100 Suppl 1(Suppl 1), S10-s17. 10.1016/j.jped.2023.08.009

Wood, J. A., & Scott, J. F. (2021). Economic impacts of chickpea grain classification: how ‘seed quality is Queen’ must be considered alongside ‘yield is King’ to provide a princely income for farmers. CROP & PASTURE SCIENCE, 72(2), 136–145. 10.1071/cp20282

Zhou, G., & Wang, Q. (2018). A new nonlinear method for calculating growing degree days. Scientific Reports, 8(1), 10149. 10.1038/s41598-018-28392-z

